# The *Arabidopsis* NOT4A E3 ligase coordinates PGR3 expression to regulate chloroplast protein translation

**DOI:** 10.1101/2020.04.02.021998

**Authors:** Mark Bailey, Aiste Ivanauskaite, Julia Grimmer, Oluwatunmise Akintewe, Adrienne C. Payne, Ross Etherington, Anne-Marie Labandera, Rory Osborne, Marjaana Rantala, Sacha Baginsky, Paula Mulo, Daniel J. Gibbs

## Abstract

Chloroplast function requires the coordinated action of nuclear- and chloroplast-derived proteins, including several hundred nuclear-encoded pentatricopeptide repeat (PPR) proteins that regulate plastid mRNA metabolism. Despite their large number and importance, regulatory mechanisms controlling PPR expression are poorly understood. Here we show that the Arabidopsis NOT4A ubiquitin-ligase positively regulates PROTON GRADIENT 3 (PGR3), a PPR protein required for translating 30S ribosome subunits and several thylakoid-localised photosynthetic components within chloroplasts. Loss of NOT4A function leads to a strong depletion of plastid ribosomes, which reduces mRNA translation and negatively impacts photosynthetic capacity, causing pale-yellow and slow-growth phenotypes. Quantitative transcriptome and proteome analyses reveal that these defects are due to a lack of PGR3 expression in *not4a*, and we show that normal plastid function is restored through transgenic PGR3 expression. Our work identifies NOT4A as crucial for ensuring robust photosynthetic function during development and stress-response, through modulating PGR3 levels to coordinate chloroplast protein synthesis.

## Introduction

The synthesis of energy from the sun, photosynthesis, supports organic life on earth. Light harvesting in green plants takes place within the specialized chloroplast organelle, believed to have arisen from engulfment of a photosynthetic prokaryote by an ancestral eukaryotic cell (Archibald, 2015). Coevolution and merging of these organisms has resulted in nuclear and chloroplast genomes separated within cellular compartments. In land plants, the chloroplast genome comprises of ∼130 genes, yet chloroplasts contain around 3000 different proteins (Zoschke and Bock, 2018). Consequently, chloroplast function requires expression not only of chloroplast encoded proteins, but a multitude of nuclear encoded genes, which are imported into chloroplasts post-translationally. One such group of nuclear derived factors is the pentatricopeptide repeat domain (PPR) containing proteins. The PPR protein family has significantly expanded in plants (∼450 in Arabidopsis, vs <10 in humans and yeast; (Schmitz-Linneweber and Small, 2008)), and members are characterized by a 35-amino acid repeat sequence that facilitates RNA binding and enables them to provide critical gene expression control within chloroplasts and mitochondria (Barkan and Small, 2014). Through binding to organellar RNAs, PPR proteins stabilize gene transcripts, facilitate post-transcriptional processing and promote translation of the encoded proteins (Barkan and Small, 2014; Manna, 2015). Whilst their function in the regulation of gene expression control within organelles has been described, including many of the RNA species to which they bind, little is known about how their expression is regulated prior to import.

Precise, selective removal of proteins is essential to cellular development and response. In eukaryotes, proteins can be marked for degradation by the megacomplex protease known as the 26S proteasome, following enzymatic attachment of a chain of ubiquitin molecules (Sadanandom et al., 2012). Ubiquitin attachment requires sequential enzyme activities: initial processing of pre-ubiquitin by deubiquitinating enzymes (DUBs), followed by bonding to an E1 activating enzyme, transfer to an E2 conjugating enzyme, and finally, conjugation to a substrate mediated by an E3 ligase enzyme (Callis, 2014). Despite the absence of the ubiquitin proteasome system (UPS) within plastids, three ubiquitin E3 ligases mediating chloroplast proteostasis have been described. The Hsc70-interacting protein (CHIP) E3 ligase was shown to target pre-plastid proteins for degradation in a chloroplast import-defective mutant background, indicating that it is required for ensuring correct and complete targeting of proteins to this organelle (Lee et al., 2009; Shen et al., 2007a; 2007b). A second E3 ligase, Plant U-box ubiquitin ligase PUB4, regulates the degradation of oxidatively damaged chloroplasts (Chlorophagy), via ubiquitination of envelope proteins (Woodson et al., 2015). Finally, a system for chloroplast associated protein degradation (CHLORAD) targets damaged components of the chloroplast transmembrane protein import machinery (TOC) (Ling et al., 2019). CHLORAD-targeted TOC subunits are ubiquitinated by the integral chloroplast outer membrane ubiquitin ligase SP1, promoting removal and delivery to the 26S proteasome via the mp85-type beta-barrel channel SP2 and AAA+ chaperone CDC48 (Ling and Jarvis, 2015; Ling et al., 2019; 2012). However, despite the evolutionary expansion and significance of the UPS in plants (accounting for more than 5% of the genome in Arabidopsis (Smalle and Vierstra, 2004)) the full extent to which the UPS can influence chloroplast function remains to be determined.

The E3 ubiquitin-ligase NOT4 contains a unique combination of RING finger and RNA Recognition Motif (RRM) domains, which places its function at the interface of proteolysis and RNA biology (Cano et al., 2010; Chen et al., 2018; Wu et al., 2018). Studies of NOT4 in yeast and animals have revealed that it associates with ribosomes, and plays a role in co-translational quality control of mRNA and protein, to ensure efficient and correct translation of polypeptides (Dimitrova et al., 2009; Duttler et al., 2013; Halter et al., 2014; Preissler et al., 2015; Wu et al., 2018). Furthermore, NOT4 plays a role in the assembly and integrity of functional proteasomes in yeast (Panasenko and Collart, 2011). Although it can exist as a monomer, NOT4 contributes to global RNA metabolism and gene expression control as a member of the CCR4-NOT complex (Albert et al., 2002; Collart and Panasenko, 2012). Whilst the NOT4 CCR4-NOT association is strong in yeast, structural and biochemical studies indicate it is more labile in animals, suggesting that, whilst general functions in translational regulation are conserved, there has been kingdom specific divergence in mechanism (Bhaskar et al., 2015; Keskeny et al., 2019). NOT4 has also been proposed to function as an N-recognin E3 ligase of the acetylation-dependent N-end rule pathway in yeast (Shemorry et al., 2013). Despite a central role for yeast and mammalian NOT4 in co-translational control and moderating the stabilities of mRNA and proteins, the presence and functions of NOT4-like E3 ligases in plants has not been investigated in detail previously.

Here we identify three NOT4-like E3 ligases in *Arabidopsis thaliana*, and show that one of these – NOT4A - has diverged toward a key role in regulating chloroplast protein biogenesis and photosynthetic function. Using genetic, biochemical, transcriptomic and proteomic approaches, we show that NOT4A is required for expression of the PPR protein PGR3, a nuclear encoded factor that is imported into plastids to promote ribosome formation and ensure efficient chloroplast-encoded protein synthesis. *not4a* mutants, which have severely depleted *PGR3* levels, share the molecular consequences of null *pgr3* mutants, having reduced abundance of plastid RNAs that are normally targeted by PGR3, and a depletion of chloroplast ribosomes. This results in concomitant defects in chloroplast protein translation, and reduced levels of many photosynthetic complexes (including Cyt b_6_f), which leads to a pale-yellow phenotype, high-light stress sensitivity and compromised photosynthetic capacity. Site directed mutational analysis of NOT4A reveals its E3 ligase and RNA binding functions are both essential to its chloroplast related function, and transgenic expression of PGR3 in *not4*a is sufficient to restore wild type-like growth and development. Our work identifies NOT4A as an important mediator of PPR controlled plastid protein biogenesis, through promotion of *PGR3* expression, uncovering a new layer of homeostatic and stress-responsive regulation controlling chloroplast form and function.

## Results

### Identification of NOT4-like protein in Arabidopsis

To identify NOT4-like proteins in plants we searched for protein sequences with homology to *Sc*NOT4 from *Saccharomyces cerevisiae*. In contrast to the single NOT4 gene present in humans, Drosophila and yeast, we identified three putative homologues in the model plant *Arabidopsis thaliana*, which we confirmed with reciprocal BLASTs. The three homologues, which we named NOT4A, -B and -C, possess 38-39% identity with *Sc*NOT4 (Figures 1A and S1), and crucially share high identity across the unique combination of RING, RRM and C3H1 domains. We isolated homozygous T-DNA insertion lines for *NOT4A-C* from publicly available collections (GABI-KAT, SALK, SAIL) and confirmed T-DNA inserts and full-length mRNA knockout (Figures S2A and B). The *not4a* line displayed a pale-yellow phenotype and a clear delay in development, flowering significantly later than wild type under normal growth conditions (Figures 1B and D). Moreover, *not4a* had a significant reduction of chlorophyll (Figure 1E), and Lugol’s staining revealed reduced starch accumulation relative to Col-0 (Figure 1F). In contrast, *not4b* and *not4c* displayed no obvious growth phenotypes, and so we decided to focus our attention on NOT4A.

**Figure 1.**
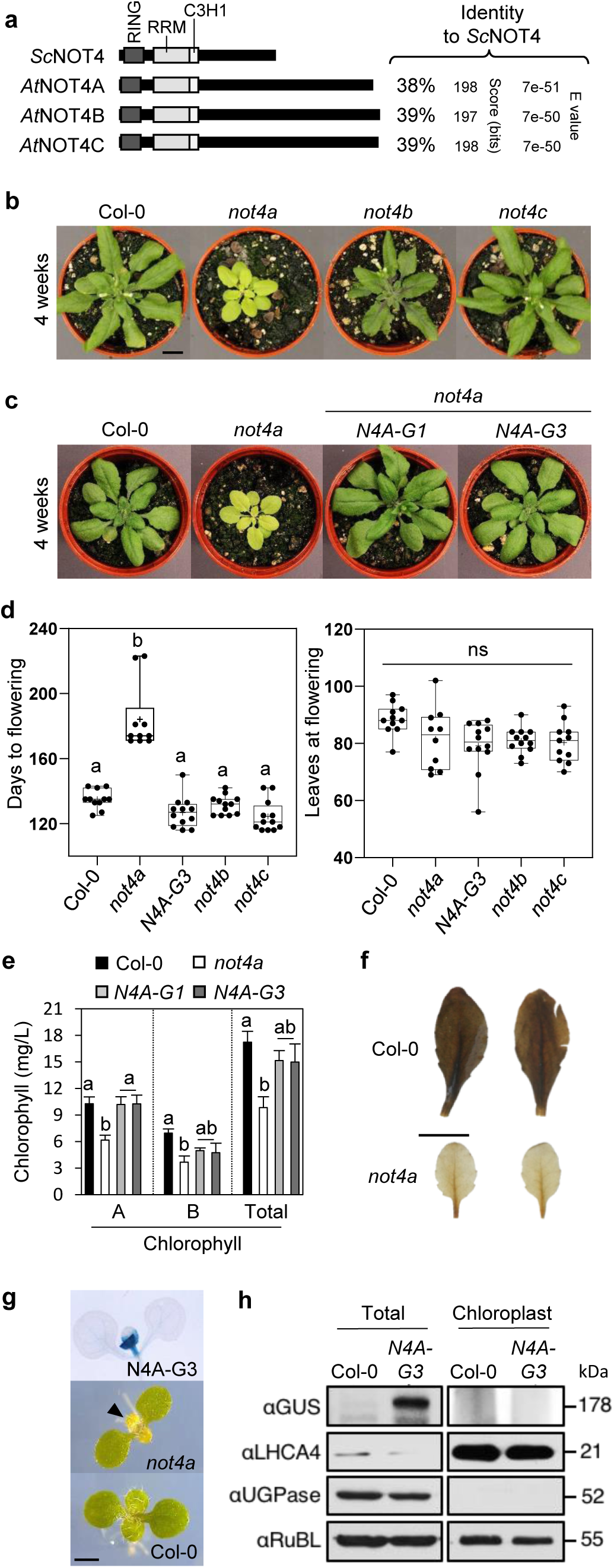
Identification of NOT4-like genes in plants. **(a)** Schematic diagram of protein domain structure and % amino acid identity for *Sachharomyces cerevisiae* (*Sc*) and *Arabidopsis thaliana* (At) NOT4 proteins. For full sequence alignments see Supplementary Figure 1. **(b)** and **(c)** Representative rosette images of 4-week old col-0, *not4a-c* mutants, and two independent *not4a* complementation (*N4A-G1* and *G3*) lines grown under long day (LD) conditions. Bar = 1cm. **(d)** Days to flowering and rosette leaf number at flowering for genotypes shown under short day (SD) conditions (n=10-12 per genotype). Box and whiskers plots show max and min, 25^th^ to 75^th^ percentiles, median and mean (+). Letters indicate one way ANOVA; Tukey’s test (p<0.01). **(e)** Chlorophyll content (A, B and total) of SD grown rosette leaves from Col-0, *not4a* and two independent complementation lines (*N4A-G1* and *G3*). Error bars = SEM. **(f)** Lugol’s iodine staining of Col-0 and *not4a* rosette leaves. Bar = 1cm. **(g)** Histochemical staining of 7-day-old seedling of the *N4A-G3* complementation line showing localisation of *pNOT4::NOT4-GUS* to the first true leaves. This correlates with where the pale-yellow phenotype first presents in *not4a* (arrowhead) relative to Col-0. Bar = 200µm. **(h)** Detection of *pNOT4::NOT4A-GUS* by anti-GUS western blot in total vs chloroplast-specific protein extracts. LHCA4 is a chloroplast enriched protein control. UGPase is a cytosol control showing efficacy of chloroplast enrichment. RuBL was used as a loading control.

To confirm the pale-yellow *not4a* phenotype was due to loss of NOT4A expression we complemented the mutant by reintroduction of full-length genomic NOT4A with a c-terminal GUS tag, driven by ∼2kb of its endogenous promoter. Two independent transgenic lines expressing the full length NOT4A protein (N4A-G1 and N4A-G3, Figure S2C), displayed wild-type greenness and normal development (Figures 1C, D and E), and NOT4A expression, determined by GUS staining, was localized to the first true leaves of seedlings where the pale mutant phenotype first presents (Figure 1G). Analysis of the full length NOT4A amino acid sequence using TargetP (Emanuelsson et al., 2007) revealed no obvious chloroplast transit peptide (cTP). In accordance with this prediction, anti-GUS western blotting of total vs chloroplast-specific protein extracts indeed revealed that NOT4A is not associated with this organelle, which is also in line with the absence of the ubiquitin proteasome system in plastids (Figure 1H).

### NOT4A is required for chloroplast and photosynthetic function

The growth, development, and starch depletion phenotypes of *not4a* point to general defects in photosynthesis. To investigate this further, we carried out an RNA seq analysis on 10-day-old *not4a* and Col-0 seedlings. This revealed a large number of differentially expressed genes (DEGs) in the mutant relative to WT (>2300 genes, P<0.05), with GO-analysis showing that a large proportion of these are chloroplast related, and implicated across all structures of this organelle (Figures 2A, B and S3, Data file 1). Carbon assimilation in stroma is undertaken by an array of higher order enzymatic complexes present within the internal membrane system of chloroplasts (thylakoids), which includes many nuclear-encoded components that were mis-regulated in the *not4a* transcriptome (Figure S3, Data file 1)(Allen et al., 2011). We therefore analysed the composition of the protein complexes within the thylakoids of Col-0, *not4a* and N4A-G3 complementation lines by blue native gel analysis (BN-PAGE) (Aro et al., 2005; Järvi et al., 2011; Rantala et al., 2017). This revealed in *not4a* a severe decrease in the accumulation of protein bands corresponding to the photosystem II monomer and Cytochrome b_6_f complex (PSII m/ Cyt b_6_f), as well as the photosystem I - NAD(P)H dehydrogenase megacomplex (PSI-NDH), when equal protein was loaded (Figure 2C and S4A). Individual subunits were then resolved by a second denaturing dimension, which confirmed lower abundance of the PetA, -B, -C and -D Cytochrome b_6_f complex subunits (Figure 2D), as well as the CP47, CP43, D2 and D1 subunits of the PSII monomer (Figure S4B). In addition, dual PAM measurements revealed a clear reduction in electron transfer rate (ETR) in *not4a*, consistent with depletion of Cytb_6_f within the thylakoid membranes (Figure 2E) (Tikhonov, 2014).

**Figure 2.**
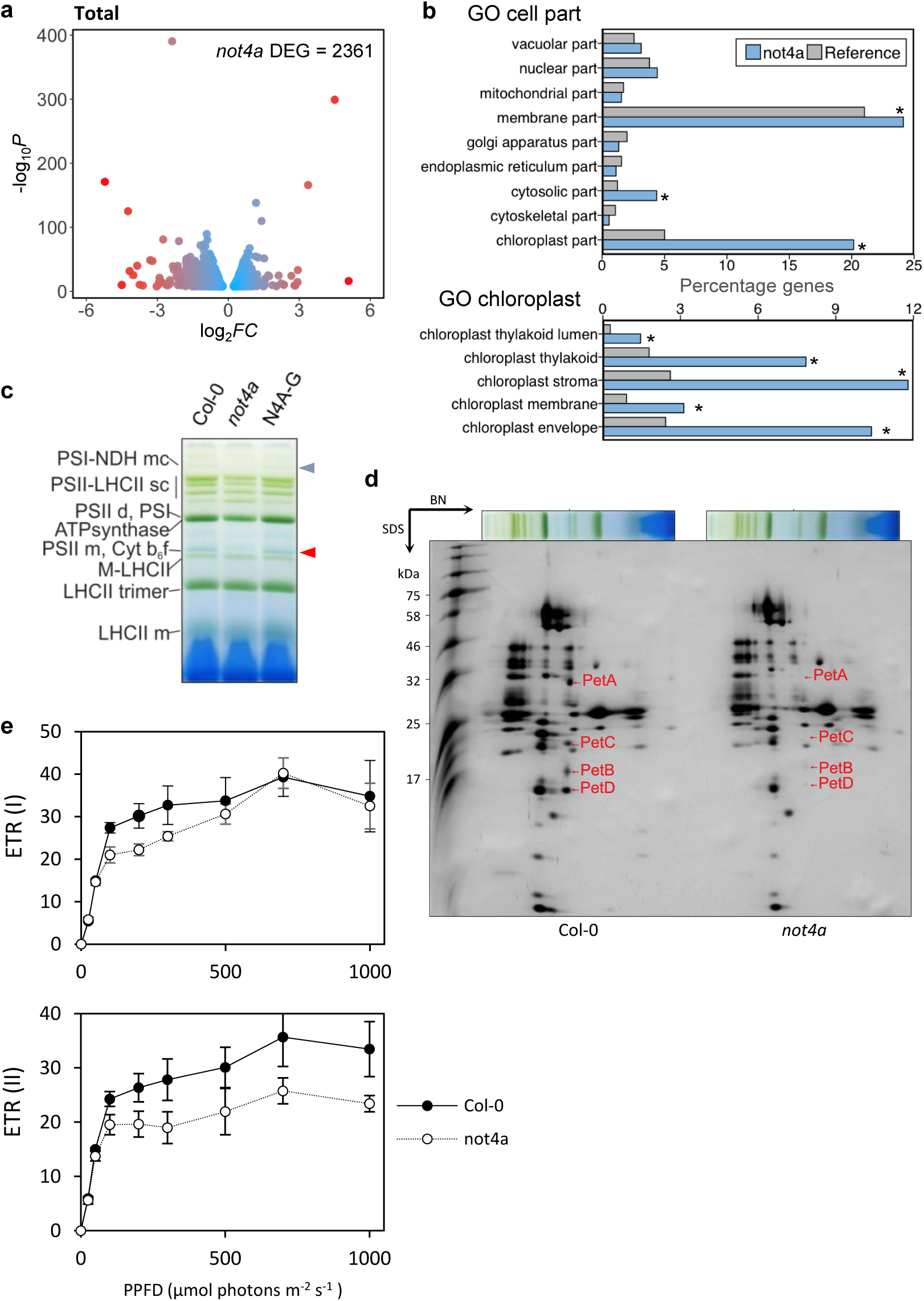
NOT4A is required for chloroplast and photosynthetic function. **(a)** Volcano plot of up and down DEGs in *not4a* vs WT. **(b)** Graphical representation of ‘cell part’ and ‘chloroplast’ Gene Ontology (GO) enrichment of DEGs in *not4a* vs WT. Asterisk refers to significant enrichment. **(c)** Blue native (BN) gel of thylakoid protein complexes from Col-0, *not4a* and N4A-G3. Plants were grown under standard conditions (100 µmol photons m^-2^ s^-1^, 8 h/16 h light/dark), and the entire thylakoid network was solubilized with 1% (w/v) dodecyl maltoside (DM) and separated using BN gel electrophoresis. 50 µg of total protein was loaded. Red and blue arrows refer to PSII m/Cyt b_6_f and PSI-NDH mc bands, respectively, which are significantly depleted in *not4a*. See also Supplemental Figure 4a. **(d)** Second dimension separation of DM-solubilised thylakoid proteins from Col-0 and *not4a* showing depletion of Cyt b_6_f components PetA, PetB, PetC and PetD in *not4a*. The protein bands were identified based on Aro et al. 2005. **(e)** Light-intensity dependence of ETR(I) and ETR(II). Dark adapted Col-0 and *not4a* plants were subjected to illumination steps of 3 min at light intensities of 25-1000 μmol photons m^-2^ s^-1^ followed by a saturating flash (700 ms) to determine F_M’_ and P_M’_. The relative rates of electron transfer through PSI and PSII, ETR(I) and ETR (II), respectively, were calculated as Y(I) x PPFD x 0.84 × 0.5 and Y(II) x PPFD x 0.84 × 0.5. Each point represents the average ± SD (n = 5).

To complement our RNA seq analysis and investigate differences in protein abundance in *not4a* vs WT in more detail, we carried out a quantitative proteomics analysis of total protein extracts as previously described (Helm et al., 2014). Overall, a substantial number of proteins were significantly changed in abundance in the mutant compared to wildtype (Figure 3A, Data file 2). Analysis of the chloroplast-specific proteins in these datasets (619 in WT, 689 in *not4a;* Figure 3B) confirmed reduced levels of the majority of the subunits making up the Cytb_6_f and NDH complexes, but remarkably also identified a broad depletion of many of the central enzymes required for photosynthesis (Figures 3C and S4C, Data file 2), which was likely masked in the blue native gels due to thylakoid enrichment and equal protein loading.

**Figure 3.**
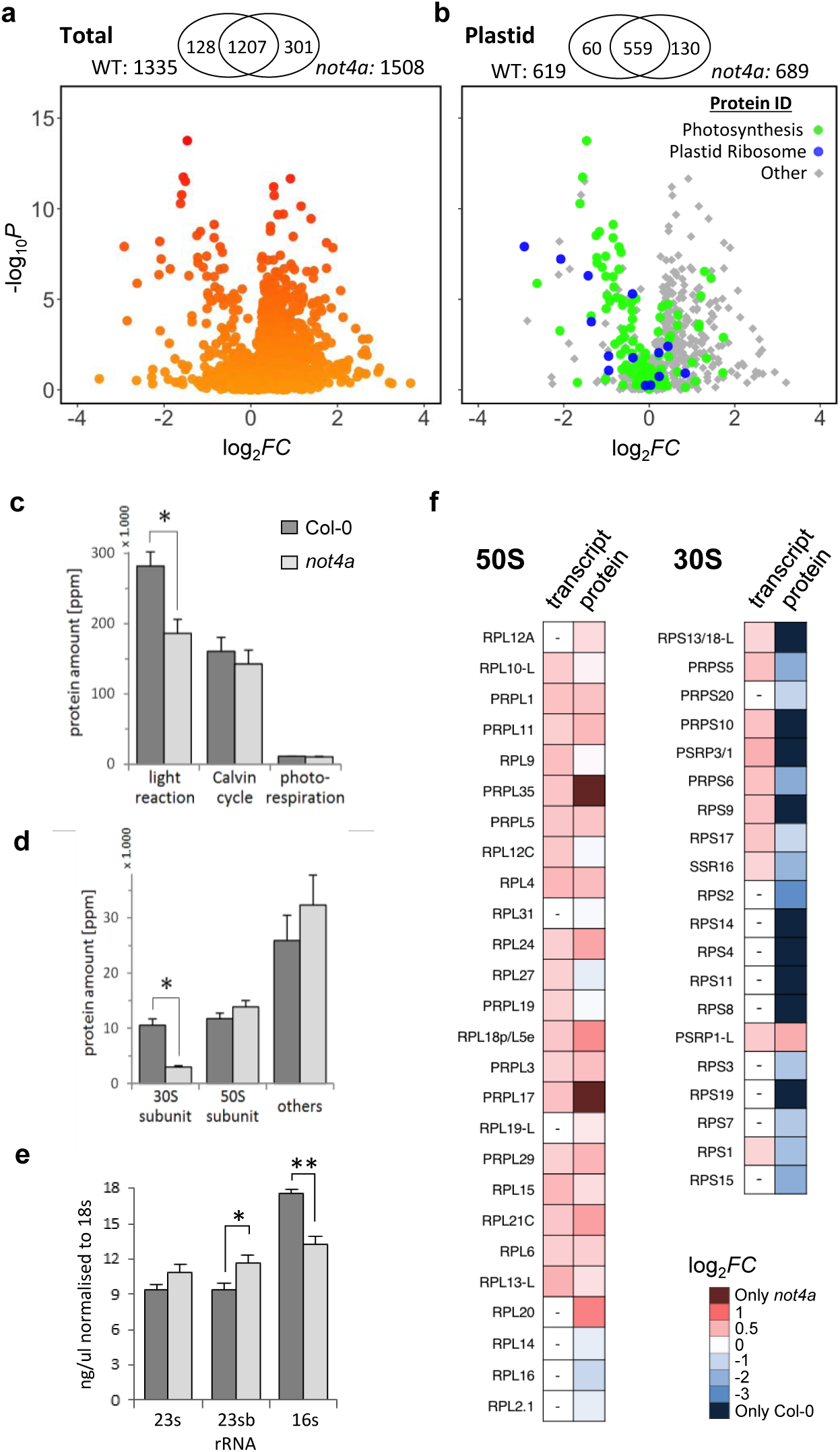
NOT4A is required for plastid ribosome biogenesis. **(a)** and **(b)** Volcano plots showing differential abundance of total and plastid-localised proteins in *not4a* vs wild type (Col-0): x-axis, log2-Fold Change values of protein abundances in *not4a* versus wildtype; y-axis, log10-p values from a two-sided t-test. For the plastid-specific graph, photosynthetic and plastid ribosome proteins are highlighted in green and blue, respectively. The number of quantified proteins and their overlap are shown in Venn diagrams above each plot. **(c)** and **(d)** Histograms of summed protein amounts (ppm) of MapMan (Thimm et al., 2004) annotated functional groups. Asterisks highlight functional groups that are significantly depleted in *not4a* vs Col-0. Error bars= SEM. **(e)** Relative amounts (ng/µl) of 50S (23s and 23sb) and 30S (16s) plastid rRNAs in *not4a* vs Col-0. Values are quantified from Tapestation data shown in Supplemental Figure 4 following normalisation to cytosolic 18s rRNA. **(f)** Heatmap showing relative transcript and protein fold-changes for 50S and 30S plastid ribosome subunits in *not4a* vs Col-0.

### NOT4A is required for plastid ribosome biogenesis

In addition to a reduction of photosynthetic proteins, we also observed a significant downregulation or absence of plastid ribosome proteins in *not4a* (Figure 3B, D and F, Data file 2). Remarkably, this effect was specific to proteins making up the 30S subunit of the chloroplast ribosome, whilst components of the 50S subunit were detected at similar or slightly increased levels relative to wild-type (Figure 3D and F). Of 20 30S subunits identified in the proteomic analysis, 18 were significantly down regulated (>1.5 fold) in *not4a* vs WT, whilst none of the 26 50S subunits identified were reduced in abundance. In support of this, we also observed a significant reduction of chloroplast 30S-associated 16S rRNA in *not4a* relative to WT, whilst levels of the 50S-associated 23S and 23Sb rRNAs were similar (Figures 3E and S5). Interestingly, the reduction of 30S subunits was only apparent at the protein level, as transcripts of the nuclear encoded subunits were in fact elevated in the RNA seq data (Figure 3F, Data file 2). This indicates that decreased 30S ribosome abundance in *not4a* is linked to a translational or post-translational, rather than transcriptional, defect.

### Plastid mRNA translation is compromised in the *not4a* mutant

The severe reduction of 30S ribosome subunits and many chloroplast encoded proteins in *not4a* (Figures 3D, F and S4C) prompted us to investigate chloroplast translation in this mutant. We observed extreme hypersensitivity of *not4a* to the chloroplast ribosome-specific inhibitor lincomycin (Figure 4A and B). Lincomycin treated Col-0 seedlings were smaller in size and more yellow than control plants, resembling untreated *not4a* mutants, corroborating the observation that chloroplast ribosome activity is perturbed in *not4a* (Figure 4A and B). In contrast, no obvious differences in the sensitivity to the cytosolic ribosome inhibitor cycloheximide (CHX) were observed (Figure S6A). Next we assessed chloroplast-specific protein synthesis in WT and *not4a* through assaying incorporation rates of the aminoacyl-tRNA analogue puromycin in isolated chloroplasts (Van Hoewyk, 2016). Puromycin labelling of nascent proteins was strongly reduced in *not4a* chloroplasts relative to WT, indicating reduced translation capacity in the mutant (Figure 4C). Alongside the lincomycin results, this reveals that the reduced levels of 30S ribosomes in *not4a* causes defects in chloroplast mRNA translation. Since a majority of photosynthetic complexes include chloroplast-encoded components (Allen et al., 2011), reduced chloroplast translation likely explains the broad reduction in photosynthetic proteins in *not4a*, with subunit imbalance leading to complex collapse i.e. degradation of unassembled nuclear subunits also (Choquet and Wollman, 2002; Peng et al., 2009, Juszkiewicz and Hegde, 2018).

**Figure 4.**
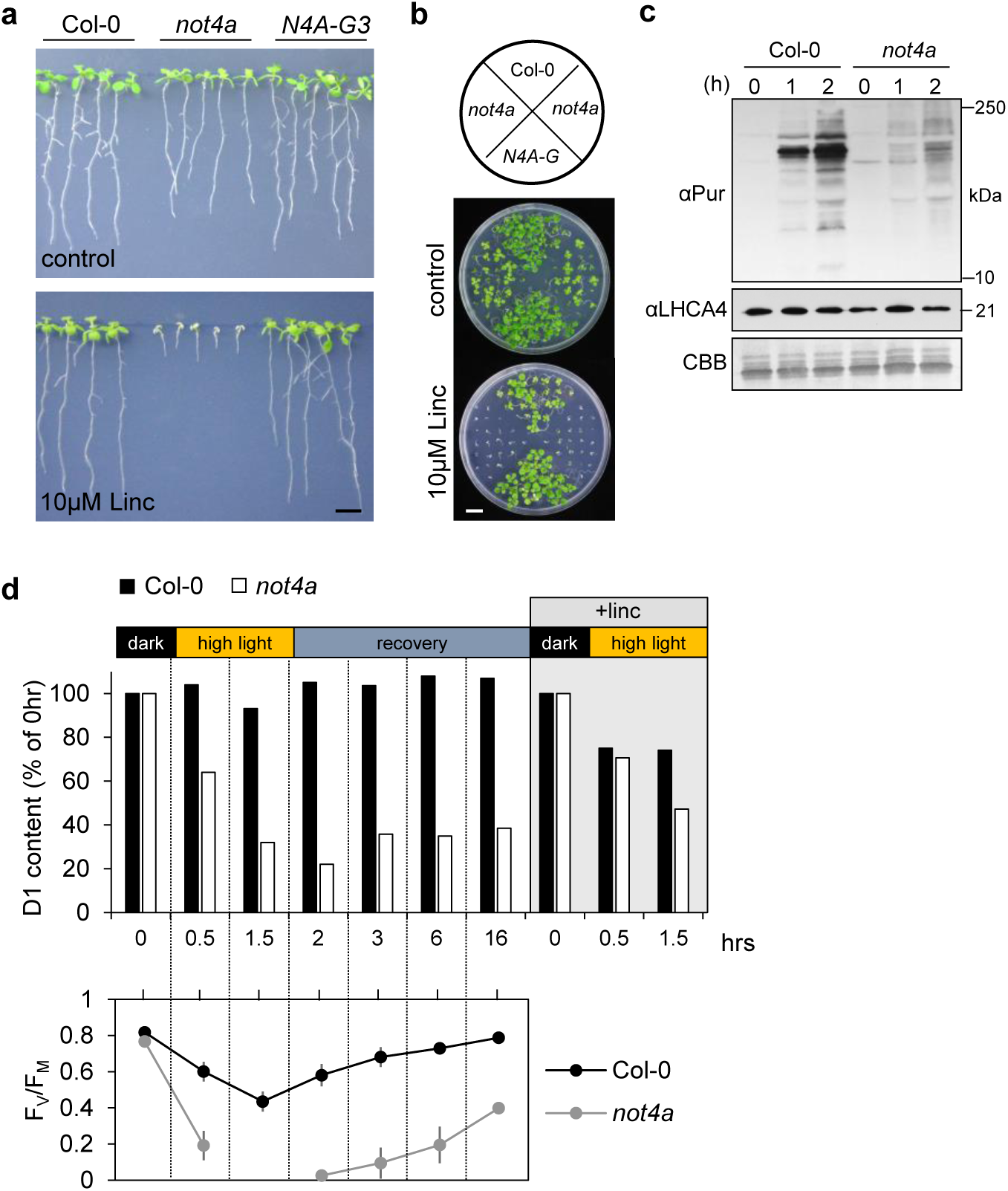
Plastid mRNA translation is compromised in the *not4a* mutant. **(a)** 10-day old WT, *not4a* and N4A-G3 complementation line seedlings grown on vertical plates +/- 10µM lincomycin. Bar = 1 cm. **(b)** same as in **(a)** but on horizontal plates. Bar = 1cm. **(c)** Protein synthesis rates in Col-0 and *not4a* chloroplasts, determined by anti-puromycin (α-Pur) western blot following a 2-hour (h) puromycin treatment time course. LHCA4 and CBB (Coomassie Brilliant blue) are shown as loading controls. **(d)** The amount of D1 protein (% of 0 hr) and PSII efficiency (F_V_/F_M_) during a photoinhibitory treatment at 1000 μmol photons m^-2^ s^-1^ with and without lincomycin (linc) and during recovery without linc at 100 μmol photons m^- 2^ s^-1^. Total leaf protein extracts were separated on an SDS-PAGE and immunoblotted with a D1-specific antibody. Representative results of three biological replicates are presented in the figure. For PSII efficiency the average ± SD (n = 5) is presented.

Reduction in *not4a* of the PSII monomer (Figures 2C, S4B and C), a repair intermediate of the bioactive PSII-LHCII super complex, suggests defects in PSII repair. The central scaffold of PSII is the chloroplast encoded D1 protein, a photolabile protein that is enzymatically broken down after light-induced damage (Li et al., 2018). The high turnover rate of the D1 protein requires rapid synthesis and replacement to reassemble the PSII complex, which is essential for photosynthesis. We examined D1 turnover and PSII efficiency during and after high light (HL) exposure of the *not4a* and Col-0 plants. After 1.5hrs of HL D1 abundance was reduced to ∼25% of starting levels in *not4a*, whereas levels in Col-0 were maintained above 90% (Figure 4D). This is in line with the detected PSII activities, measured as F_v_/F_M_, which decreased in both lines, but less severely in Col-0. After 16 hours of recovery in standard conditions (GL), Col-0 D1 levels returned to 100%, whilst in *not4a* they remained below 50% (Figure 4D), implying defects in D1 biosynthesis. Reduction of D1 when lincomycin was applied during HL treatment to inhibit translation, confirmed that D1 synthesis compensates for damaged and degraded protein in Col-0. Taken together these data show that NOT4A is required to maintain homeostatic and stress-responsive translational productivity in chloroplasts.

### Functional domain analysis of NOT4A

To investigate the mechanistic connection between NOT4A and chloroplast function in more detail, we mutated the conserved RING and RRM domains in *pNOT4A::NOT4A-GUS* and tested if these variants could still complement the *not4a* mutant. These mutations were based on conserved homologies to yeast *Sc*NOT4 where the activities of both domains were successfully knocked out previously (Figure 5A and S1)(Chen et al., 2018; Dimitrova et al., 2009; Mulder et al., 2007). NOT4A proteins containing either a single L11A mutation in the N-terminal RING domain, three point mutations in the RRM domain (G137A, Y166A and C208A), or all four substitutions were expressed under the endogenous promoter (Figure 5B). We observed particularly high levels of the L11A RING mutant variant, suggesting that abolishing RING activity may enhance NOT4A stability by preventing auto-ubiquitination, a common feature in E3 ubiquitin-ligase regulation (de Bie and Ciechanover, 2011). Mutation of the RRM however, resulted in reduced NOT4A levels, presumably due to disruption of protein activity leading to enhanced auto-ubiquitination, a notion supported by comparatively increased abundance of NOT4A with combined RING/RRM mutations. In contrast to complementation with WT *pNOT4A::NOT4A-GUS*, none of the mutant variants were functional *in planta*, as evidenced by the incapacity to restore WT tolerance to lincomycin or revert the pale-yellow phenotype of the *not4a* mutant (Figure 5C and D). Significantly, NOT4A expression was upregulated when seedlings were grown in the presence of lincomycin, suggesting NOT4A expression is controlled by plastid to nucleus retrograde signalling during chloroplast translational stress (Figure 5E). Overall, we can deduce that both domains of NOT4A are required for its homeostatic and stress-responsive roles in regulating chloroplast function.

**Figure 5.**
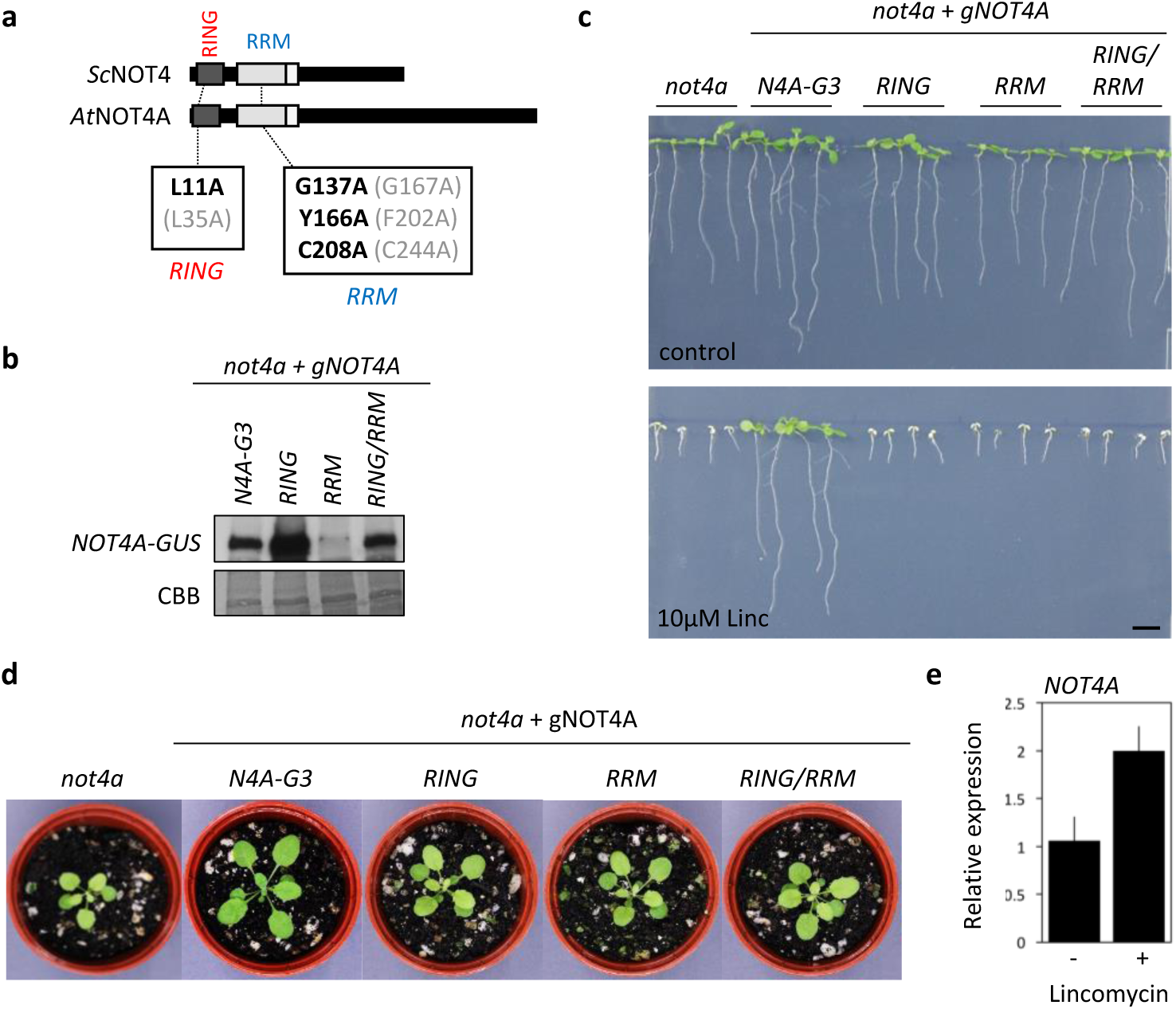
Functional domain analysis of NOT4A. **(a)** Schematic diagram showing position of point mutations in RING and RRM domain. Mutations in black are those introduced to the NOT4A, which correspond to equivalent conserved residues in ScNOT4 (shown in grey), additionally highlighted in yellow in Supplementary Figure 1 sequence alignments. **(b)** anti-GUS western blot of steady state levels of *pNOT4A::NOT4A*-GUS variants expressed in *not4a*. **(c)** 10-day old *not4a* and *pNOT4A::NOT4A*-GUS complementation lines grown on vertical plates +/- 10µM lincomycin. Bar = 1 cm. **(d)** Representative rosette images of 3-week old *not4a* and *pNOT4A::NOT4A*-GUS complementation lines under long day (LD) conditions. Bar = 1cm. **(e)** Quantitative RT-PCR (qPCR) of *NOT4A* in 14-day-old Col-0 lines grown on media +/- 10µM Lincomycin. Expression levels normalised to *ACTIN7* and shown relative to untreated Col-0. Data are average of three biological replicates. Error bars = SEM.

### The *not4a* mutant mimics the *pgr3* pentatricopeptide mutant

To identify potential causal agents of 30S ribosomal depletion in *not4a* we analysed the DEGs annotated with chloroplast functions (GO:0009507) in the *not4a* RNAseq dataset (Figure 6A, Data file 1). A total of 828 DEGs were identified, with a vast majority of these (699) being upregulated relative to WT. Amongst these genes were 34 chloroplast-targeted PPR proteins that regulate organellar gene expression, which may be upregulated to compensate for compromised translation in the mutant (Figure S6B, Data file 2) (Bryant et al., 2011; Myouga et al., 2010; 2013). Remarkably, we found that one of these chloroplast-targeted PPR proteins, Proton Gradient Regulated 3 (PGR3), was downregulated to undetectable levels in the mutant, but restored to WT levels in the N4A-G3 complementation line (Figure 6B). PGR3 was originally identified in a screen for mutants with defects in non-photochemical quenching (NPQ) determined by chlorophyll fluorescence (Yamazaki et al., 2004). The compromised NPQ and ETR of *pgr3* mutants was subsequently attributed to PGR3’s role in promoting stabilisation and translation of the chloroplast *PetL* operon (which encodes for the Cyt b_6_f subunits PetL and PetG), as well as a role in regulating production of the NDH subunit NdhA. As such, *pgr3* mutants have reduced levels of Cyt b_6_f and NDH, similar to *not4a* (Figures 2C, D and S4A, B and C) (Cai et al., 2011; Fujii et al., 2013; Rojas et al., 2018; Yamazaki et al., 2004). Moreover, *pgr3* mutants in maize were shown to have reduced levels of chloroplast ribosomes, which resembles our observations in *not4a* (Belcher et al., 2015). These similarities therefore suggested that the defects in *not4a* might be due to highly reduced levels of PGR3 in this mutant.

**Fig 6.**
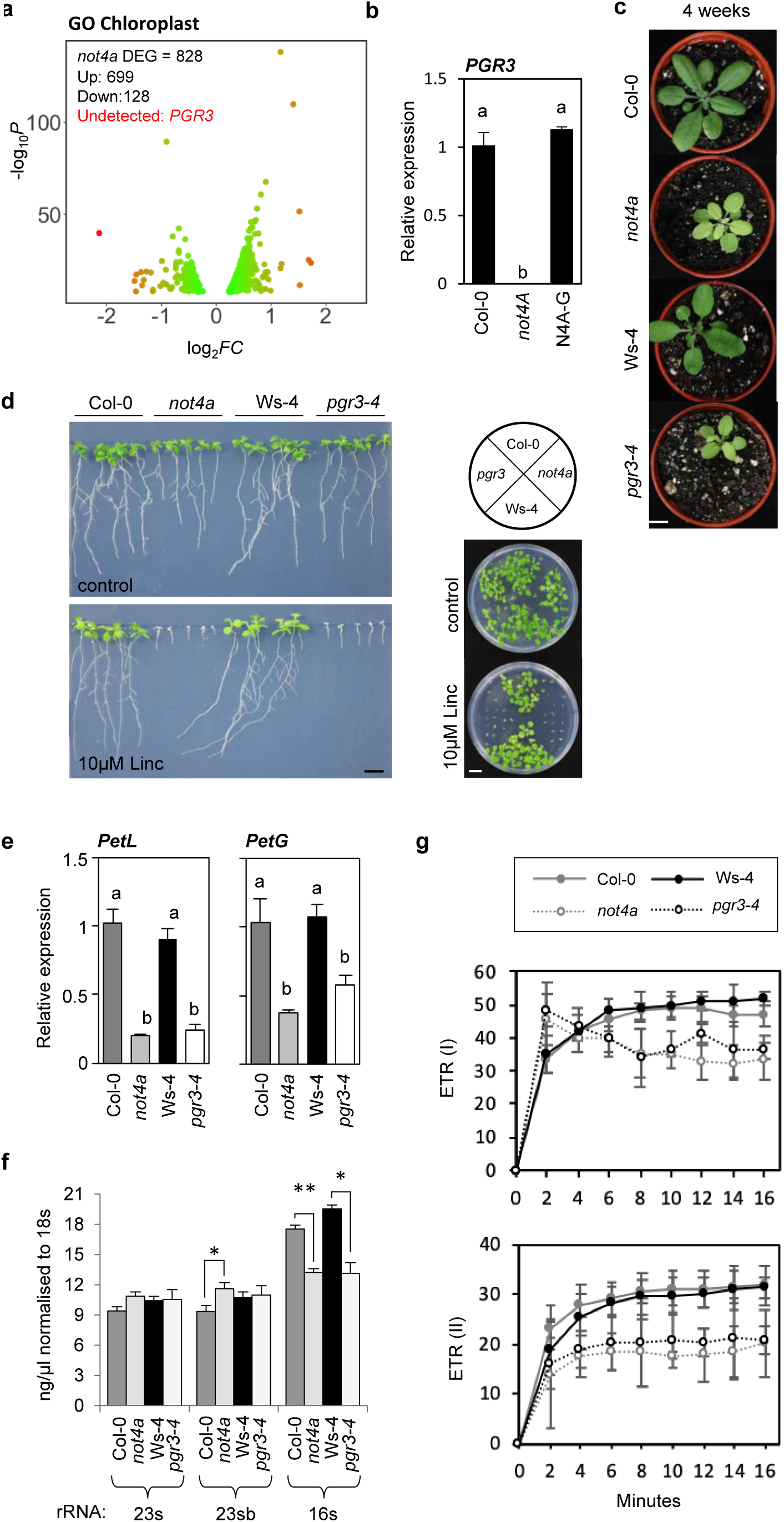
The *not4a* mutant mimics the *pgr3* pentatricopeptide mutant. **(a)** Volcano plot of GO chloroplast-associated (GO:0009507) DEGs in *not4a* vs WT. **(b)** Quantitative PCR (qPCR) of *PGR3* in Col-0, *not4a* and N4A-G3 complementation line. Expression levels normalised to *ACTIN7* and shown relative to Col-0 WT. Data are average of three biological replicates. Letters indicate one way ANOVA; Tukey’s test (p<0.01). **(c)** Representative rosette images of 4-week old Col-0, *not4a*, Ws-4 *and pgr3-4* lines grown under long day (LD) conditions. Bar = 1cm. **(d)** 10-day old Col-0, *not4a*, Ws-4 *and pgr3-4* seedlings grown on vertical and horizontal plates +/- 10µM lincomycin. Bar = 1 cm. **(e)** qPCR of *PetL and PetG* in 10-day old Col-0, *not4a*, Ws-4 *and pgr3-4* seedlings. Expression levels normalised to *ACTIN7* and shown relative to Col-0 WT. Data are average of three biological replicates. Letters indicate one way ANOVA; Tukey’s test (p<0.01). Error bars = SEM. **(f)** Relative amounts (ng/μl) of 50S (23s and 23sb) and 30S (16s) plastid rRNAs in Col-0, *not4a*, Ws-4 *and pgr3-4* seedlings. Values are quantified from Tapestation data shown in Supplemental Figure 4, following normalisation to cytosolic 18s rRNA. Col-0 and *not4a* data are same as in figure 3e. Error bars = SEM. **(g)** ETR(I) and ETR(II) during an induction curve measurement. Dark adapted Col-0, *not4a*, Ws-4 and *pgr3* plants were illuminated with 1000 μmol photons m^-2^ s^-1^ actinic light and a saturating flash was applied every 2 min to determine F_M’_ and P_M’_. The relative rates of electron transfer through PSI and PSII, ETR(I) and ETR (II), respectively, were calculated as in Figure 2. Each point represents the average ± SD (n = 3-4).

To test this further, we acquired a recently isolated null mutant of *PGR3* in the Arabidopsis Wassilewskija-4 (Ws-4) ecotype, *pgr3-4* (Rojas et al., 2018). *Pgr3-4* shares a similar delayed development and pale-yellow phenotype to *not4a* (Figure 6C). Reduced transcripts of *PetL* and *PetG* were detected in both *not4a* and *pgr3-4* relative to their respective wildtypes, consistent with the requirement for PGR3 in stabilising these mRNAs (Figures 6E and S6C). Consequently, this results in a similar reduction in ETR in both *not4a* and *pgr3-4* (Figure 6G). To determine if PGR3 is required for chloroplast translation, we tested sensitivity of the mutant to lincomycin. Here, *pgr3-4* displayed a similar degree of hypersensitivity as *not4a (*Figure 6D). Furthermore, we observed a significant reduction in 16s rRNAs in *pgr3-4*, which points to a comparable depletion of 30S subunits in both mutants (Figure 6F).

In Arabidopsis PGR3 was recently shown to control stability and stimulate expression of the 30S *Rps8* and 50S *Rpl14* ribosomal subunits (Rojas et al., 2018). Interestingly a second PPR protein, SVR7, also regulates *Rpl14* stability, but does not affect *Rps8* translation (Rojas et al 2018; Zoschke et al., 2013). We observed enhanced levels of SVR7 protein in *not4a* relative to WT (Figure S6B), perhaps compensating for loss of *PGR3*, and explaining why 50S subunits are not affected. In contrast, defects in *Rps8* translation due to loss of PGR3 in *not4a* likely explains why ribosome depletion is specific to the 30S subunit, since protein complex stoichiometries are highly regulated, with the inability to assemble complete complexes often leading to degradation of orphan subunits (Juszkiewicz and Hegde, 2018; Taggart et al., 2020). In addition, upregulation of PPR proteins SOT1, EMB2654, PPR4 and PPR2 (which all promote chloroplast rRNA maturation), and GUN1 (implicated in the production of chloroplast ribosomal subunits RPS1 and RPL11) present further evidence of compensatory responses to reduced ribosome abundance and protein synthesis within *not4a* chloroplasts (Figure S6B)(Aryamanesh et al., 2017; Lee et al., 2019; Lu et al., 2011; Tadini et al., 2016; Wu et al., 2016).

### PGR3 can rescue the chloroplast associated defects in *not4a*

Our data indicate that the *not4a* defects in chloroplast ribosome abundance, protein translation and photosynthetic function are due to loss of *PGR3* expression in this mutant. We therefore investigated if reintroducing PGR3 in *not4a* can revert these phenotypes by transforming *pPGR3::PGR3-YFP* constructs into the mutant and Col-0. PGR3-YFP protein levels were comparable across the Col-0 and *not4a* lines, whilst *PGR3* transcripts were elevated between 2-and 6-fold above WT levels in all transgenics, with *YFP-*specific qPCR corroborating the proportional pattern of expression between lines (Figures 7A, B and S7A). Interestingly, when compared to their untransformed backgrounds, two of the three Col-0 (lines 1 and 2) and all *not4a* transgenics had higher transcript levels than attributable to the double *PGR3* gene copy number expected, suggesting *pPGR3::PGR3-YFP* is uncoupled from normal endogenous control (Figure 7B).

**Fig 7.**
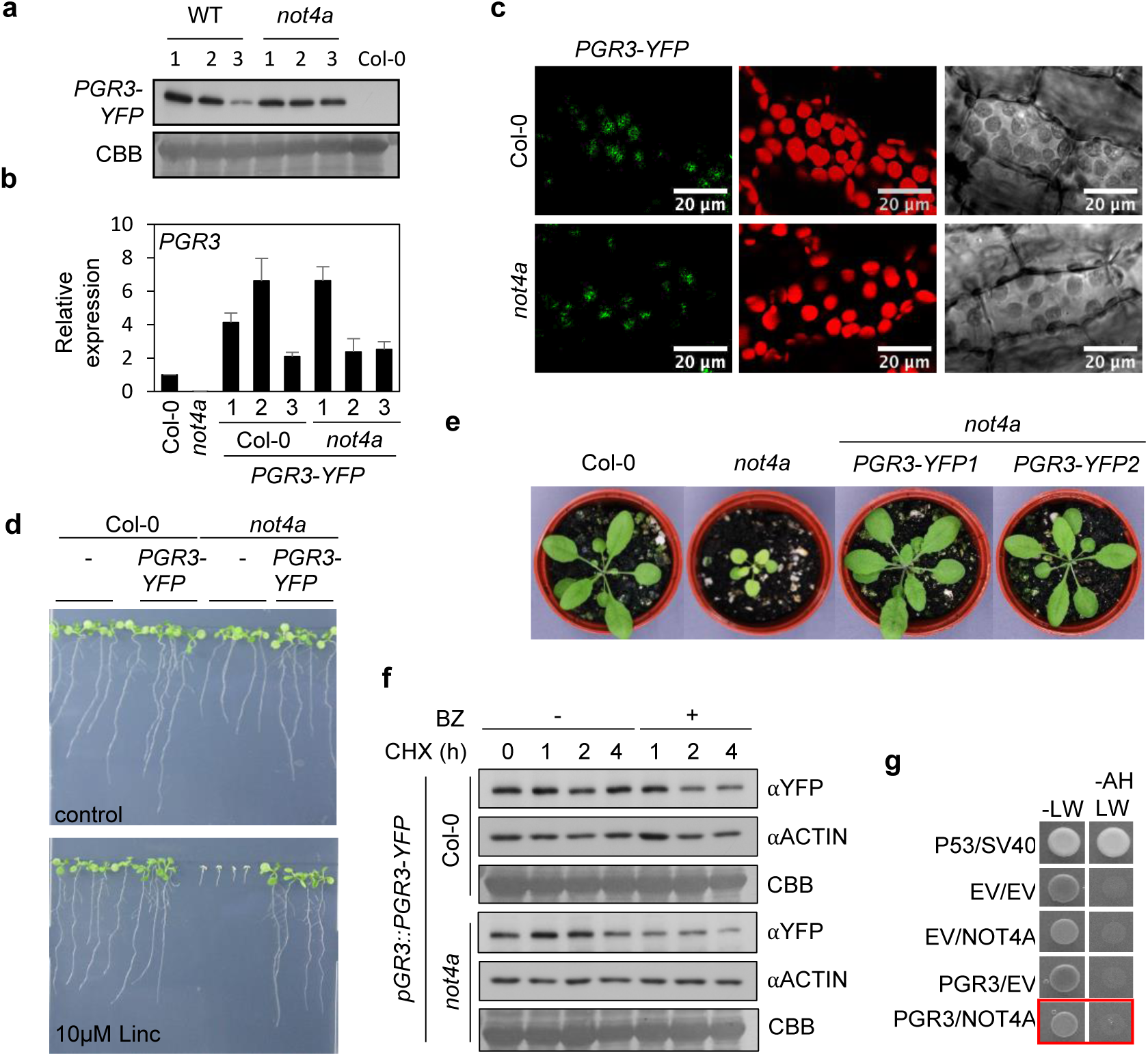
PGR3 can rescue the chloroplast associated defects in *not4a*. **(a)** Anti-YFP western blot of steady state PGR3-YFP protein levels in three independent Col-0 and *not4a* lines expressing *pPGR3::PGR3-YFP*. **(b)** Quantitative PCR (qPCR) of *PGR3* in Col-0, *not4a* and the *PGR3-YFP* lines from (a). Expression levels normalised to *ACTIN7* and shown relative to Col-0 WT. Data are average of three biological replicates. Error bars = SEM. **(c)** Confocal images of hypocotyl cells in Col-0 and *not4a* lines expressing *pPGR3::PGR3-YFP*. Panels show PGR3-YFP in green (left), chloroplasts in red (middle) and bright field view (right). **(d)** 10-day old Col-0, *not4a* and PGR3-YFP transgenic seedlings grown on vertical plates +/- 10µM lincomycin. Bar = 1 cm. **(e)** Representative rosette images of 4-week old Col-0, *not4a and not4a PGR3-YFP* lines grown under long day (LD) conditions. Bar = 1cm. **(f)** Cycloheximide (CHX) chase of PGR3-YFP and ACTIN in Col-0 and *not4a* +/- bortezomib (BZ). **(g)** Yeast two hybrid assay between NOT4A and PGR3. EV = empty vector. P53/SV40 = positive control. Growth on -LW confirms successful transformation; growth on - AHLW denotes interaction.

PGR3-YFP localized as expected within chloroplasts in both backgrounds (Figure 7C and S7B). Since NOT4 homologs function as *bona fide* E3 ubiquitin ligases to control proteasomal degradation of substrates in other organisms, and given that the NOT4A RING mutant is non-functional in Arabidopsis (Figure 5), we tested if NOT4A might target PGR3 to the UPS. Inhibition of protein synthesis using cycloheximide (CHX) did not impact upon PGR3-YFP abundance in WT or *not4a* lines, nor did proteasome inhibition with bortezomib (BZ) enhance its stability, indicating PGR3 is not a proteolytic target of NOT4A E3 ligase activity (Figure 7F). This is further supported by an absence of a protein-protein interaction between NOT4A and PGR3 when assayed by yeast-two hybrid (Figure 7G).

Remarkably however, introduction of the *pPGR3::PGR3-YFP* transgene was able to complement the pale-yellow and lincomycin-sensitive phenotypes of *not4a* (Figure 7D and E). Given that we have shown it is possible to express functional PGR3 driven by its native promoter in the *not4a* mutant, we backcrossed *not4a* to WT Col-0 and reselected for the *not4a* mutation to ensure the endogenous *PGR3* was not aberrant, and showed that the *not4a* phenotype persisted in backcrossed plants bred to homozygosity for the T-DNA insert. This, along with our prior observation that reintroduction of WT *pNOT4A::NOT4A-GUS* into *not4a* can restore *PGR3* transcript levels (Figures 1C, 4A and 6B), indicates NOT4A is required for *PGR3* expression and suggests that the inclusion of c-terminal YFP and/or a lack of regulatory elements in the reintroduced *pPGR3::PGR3* construct decouples *PGR3* regulation from NOT4A control in the mutant. This scenario is further supported by increased accumulation of *PGR3* transcripts in the Col-0 and *not4a pPGR3::PGR3* transgenic lines relative to WT (Figure 7B).

## Discussion

As essential components of the gene expression machinery in plastids, nuclear encoded PPR proteins must be dynamically controlled in order to meet the fluctuating demands of the light-harvesting chloroplast organelle. However, comparatively little is known about how PPR proteins are regulated. Here we show that expression of PGR3, a large PPR protein that regulates chloroplast ribosome biogenesis through promoting the translation of core chloroplast-encoded 30S subunits, is regulated by the cytosolic E3 ubiquitin ligase NOT4A.

We found that *not4a* mutants have photosynthetic and growth defects that are a consequence of reduced plastid ribosome biogenesis. Transcripts encoding PGR3 are undetectable in *not4a*, and *pgr3* mutants share functional defects in chloroplast activity. Remarkably, *not4a* chloroplast function was restored by introducing PGR3-YFP expressed from the endogenous PGR3 promoter sequence. The transgene led to higher *PGR3* transcript abundance in both Col-0 and *not4a* lines relative to that of WT plants. Based on known activities of NOT4 in yeast and mammals, we postulate that NOT4A is required to suppress negative regulation of *PGR3* expression at regulatory sequences outside of the reintroduced PGR3-encoding transgene, such as the 3’ UTR or distal elements, and that this regulation may act to integrate and transduce chloroplast stress signals to fine-tune plastid protein biosynthesis through modulating PGR3 abundance.

NOT4-like proteins are unique amongst E3 ubiquitin-ligases, in that they consist of both a RING domain an RNA binding RRM domain (Cano et al., 2010). We showed that both of these domains in NOT4A are required for PGR3 expression, but the molecular connection between NOT4A and PGR3 is yet to be defined. Whilst the repetitive domain structures of the PPR proteins enable precise and selective RNA binding, they may present a translational challenge to ribosomes. Two profiling studies of the Arabidopsis RNA degradome taken together, identified five PPR genes as targets for co-translational mRNA decay, with further analysis identifying three nucleotide periodicity within one PRR transcript suggesting ribosome stalling (Hou et al., 2016; Yu et al., 2016). PGR3 is the second largest and most structurally repetitive (containing 25 repeats, determined by RADAR (Madeira et al., 2019)) of the 34 DEG chloroplast targeted PPR genes in *not4a*. Significantly, Mammalian NOT4 was found to bind stalled ribosome complexes undertaking co-translational import into damaged mitochondria, initiating quality control, and ultimately mitophagy (Wu et al., 2018). Hence, potential co-translational regulation of *PGR3* transcripts by NOT4A warrants further investigation in plants.

In other organisms NOT4 contributes to post-transcriptional control as a component of the CCR4 NOT complex, central to 3’ deadenylation and mRNA decay, and in Arabidopsis, RNA binding activities of NOT4B and NOT4C were previously observed in a proteome-wide survey of RNA-binding proteins (Marondedze et al., 2016). Although the presence of the CCR4-NOT complex has yet to be established in plants, a recent proteomic characterisation of Target of rapamycin (TOR) signalling in Arabidopsis identified NOT4A as a TOR target, and NOT4B as an LST8 interactor along with other conserved CCR4-NOT core subunits, suggesting the plant NOT4s may indeed be implicated in mRNA decay within this complex (Van Leene et al., 2019). Furthermore, in humans and Drosophila, NOT4 contains a C-terminal sequence required for binding to the CCR4-NOT linker protein CAF40; a similar sequence was identified in Arabidopsis NOT4A (Keskeny et al., 2019). Notably, Drosophila NOT4 competes for the same CAF40 binding site as 3’UTR RNA binding proteins, BAG of MARBLES (BAM) and Roquin-1 (Keskeny et al., 2019; Sgromo et al., 2018; 2017). If functional BAM or Roquin-1 homologues exist in plants, NOT4A may inhibit the 3’UTR recruitment of *PGR3* to mRNA decay in plants. Exploring the *in vivo* targets of E3 ligase and RNA binding activities of all three Arabidopsis NOT4 proteins, in addition to determining their association and function within a putative CCR4 NOT complex, will help to shed light not only on how NOT4A regulates PGR3 expression, but also how these enigmatic E3 ubiquitin ligases influence other aspects of plant biology.

## Materials and methods

### Arabidopsis growth and transgenic lines

Seed were sown on compost mixed to a ratio 4:2:1 of Levington F2 compost, vermiculite and perlite, or sterile half strength Murashige & Skoog (MS) medium with 0.8% agar made with purified water and autoclaved for 15 minutes at 121°C. Plants were grown in Weiss Technik fitotron SGC 120 biological chambers with 16 hours light/ 8 hours dark cycles at 22°C (long day), or 12 hours light/ 12 hours dark at 22°C (short day).

T-DNA insertion mutants were identified from the GABI-KAT *not4a* (GABI_134E03)(Kleinboelting et al., 2012), SALK *not4b* (SALK_079194)(Alonso et al., 2003), SAIL *not4c* (SAIL_274_D03)(Sessions et al., 2002), collections from the Nottingham Arabidopsis Stock Centre (NASC) and the *pgr3-4* (line FLAG 086D06) from the Versailles *Arabidopsis thaliana* Stock Center (Rojas et al., 2018). Homozygous T-DNA insertions were identified by PCR with primers designed by T-DNA express and null expression confirmed by RT-PCR (Figure S2A and B).

Arabidopsis lines were transformed with the genomic sequences of *NOT4A* and *PGR3* plus ∼2KB of the upstream promoters cloned into Invitrogen pENTR™/D-TOPO™ (ThermoFisher-K240020) (For primer sequences see Data sheet 3) and sequenced before ligations into pGWB533 and pGWB540 constructs respectively (Nakagawa et al., 2007), using the *Agrobacterium tumefaciens* (GV3101 pMP90) floral dip method (Zhang et al., 2006).

### Histochemical staining

GUS staining was performed using 1% potassium ferricyanide and potassium ferrocyanide added to the GUS stain solution (0.1M PBS, pH 7.0, 2mM X-gluc and Triton-X-100 (0.1% v/v) **(Jefferson et al., 1987).** Seedlings were submerged in 1 ml of GUS stain solution and were incubated at 37°C in the dark for 24 hours. GUS stain solution was replaced by 1 ml of fixative (3:1 ethanol:acetic acid, 1% Tween v/v). Samples were incubated at room temperature with gentle shaking and fixative refreshed until tissues appeared cleared. Cleared samples were then mounted onto microscope slides in 50% glycerol and images captured with a bifocal light microscope.

Starch content was assessed using Lugol’s iodine staining of 6 week old short day grown rosette leaves as described (Tsai et al., 2009). Tissue was pre-cleared with ethanol and washed in distilled water before adding Lugol’s solution with rocking at room temperature (6mM iodine, 43mM KI, and 0.2M HCl), this was then washed with distilled water until clear.

### Phenotypic assays

Flowering time determined from 12 plants per line, grown under short and long day conditions. Plants were assessed daily, once a bolt of >1cm was produced the number of rosette leaves and day number was recorded.

Lincomycin sensitivity was assayed on half MS agar plates supplemented with 10µM. Lincomycin hydrochloride monohydrate (VWR-ALEXBML-A240). Sterilised seed were plated and stratified for 48 hours, then grown for 7-10 days with 16 hour light/ 8 hour dark cycles at 22°C.

Chlorophyll was quantified from 60mg of frozen adult leaf tissue. 1.8ml of DMF at 4°C was added and tubes inverted twice and incubated for 16 hours at 4°C. Absorbance was measured using BMG labtech FLUOstar OPTIMA spectrometer at 664.5nm 647nm for four biological replicates per line, chlorophyll content calculated as (Wellburn, 1994).

### RNA seq analysis

RNA was extracted using Qiagen Plant RNAeasy extraction kit as per manufacturers guidelines. RNA degradation and contamination was monitored on 1% agarose gels. Transcriptome sequencing and analysis was performed by Novogene. RNA purity was checked using the NanoPhotometer^®^ spectrophotometer (IMPLEN, CA, USA). RNA concentration was measured using Qubit^®^ RNA Assay Kit in Qubit® 2.0 Flurometer (Life Technologies, CA, USA). RNA integrity was assessed using the RNA Nano 6000 Assay Kit of the Bioanalyzer 2100 system (Agilent Technologies, CA, USA). A total amount of 3 μg RNA per sample was used as input material for the RNA sample preparations. Sequencing libraries were generated using NEBNext® UltraTM RNA Library Prep Kit for Illumina® (NEB, USA) following manufacturer’s recommendations. mRNA was purified from total RNA using poly-T oligo-attached magnetic beads. Fragmentation was carried out using divalent cations under elevated temperature in NEBNext First Strand Synthesis Reaction Buffer(5X). First strand cDNA was synthesized using random hexamer primer and M-MuLV Reverse Transcriptase (RNase H^-^). Second strand cDNA synthesis was subsequently performed using DNA Polymerase I and RNase H. Remaining overhangs were converted into blunt ends via exonuclease/polymerase activities. After adenylation of 3’ ends of DNA fragments, NEBNext Adaptor with hairpin loop structure were ligated to prepare for hybridization. In order to select cDNA fragments of preferentially 150∼200 bp in length, the library fragments were purified with AMPure XP system (Beckman Coulter, Beverly, USA). Then 3 μl USER Enzyme (NEB, USA) was used with size-selected, adaptor-ligated cDNA at 37°C for 15 min followed by 5 min at 95 °C before PCR. Then PCR was performed with Phusion High-Fidelity DNA polymerase, Universal PCR primers and Index (X) Primer. At last, PCR products were purified (AMPure XP system) and library quality was assessed on the Agilent Bioanalyzer 2100 system. The clustering of the index-coded samples was performed on a cBot Cluster Generation System using HiSeq PE Cluster Kit cBot-HS (Illumina) according to the manufacturer’s instructions. After cluster generation, the library preparations were sequenced on an Illumina Hiseq platform and 125 bp/150 bp paired-end reads were generated.

Raw data (raw reads) of fastq format were firstly processed through in-house perl scripts. In this step, clean data (clean reads) were obtained by removing reads containing adapter, reads containing ploy-N and low quality reads from raw data. At the same time, Q20, Q30 and GC content the clean data were calculated. All the downstream analyses were based on the clean data with high quality. Reference genome and gene model annotation files were downloaded from genome website directly. Index of the reference genome was built using Bowtie v2.2.3 and paired-end clean reads were aligned to the reference genome using TopHat v2.0.12. We selected TopHat as the mapping tool for that TopHat can generate a database of splice junctions based on the gene model annotation file and thus a better mapping result than other non-splice mapping tools. HTSeq v0.6.1 was used to count the reads numbers mapped to each gene. And then FPKM of each gene was calculated based on the length of the gene and reads count mapped to this gene. FPKM, expected number of Fragments Per Kilobase of transcript sequence per Millions base pairs sequenced, considers the effect of sequencing depth and gene length for the reads count at the same time, and is currently the most commonly used method for estimating gene expression levels (Trapnell et al., 2010). Differential expression analysis of two conditions/groups (two biological replicates per condition) was performed using the DESeq R package (1.18.0). DESeq provide statistical routines for determining differential expression in digital gene expression data using a model based on the negative binomial distribution. The resulting P-values were adjusted using the Benjamini and Hochberg’s approach for controlling the false discovery rate. Genes with an adjusted P-value <0.05 found by DESeq were assigned as differentially expressed. Gene Ontology (GO) enrichment analysis of differentially expressed genes was implemented using AgriGO (Tian et al., 2017).

### Quantitative RT-PCR

Arabidopsis seedlings were frozen in liquid nitrogen and ground to a fine powder. RNA was extracted using the Qiagen Plant RNAeasy kit as per manufacturers recommendations. RNA was quantified using a Thermo Scientific NanoDropTM 1000 Spectrophotometer. 1.5µg of RNA was treated with RQ1 DNase (Promega-M6101) as per manufacturers recommendations. cDNA was synthesized using oligo dT or random hexamers (for analysis of chloroplast encoded petL and -G transcripts) and SuperScript® II Reverse Transcriptase. cDNA was assessed for genomic DNA contamination using intron spanning primers for *ACTIN7*. Quantitative PCR primers were designed using the NCBI primer BLAST (Geer et al., 2010) and primer annealing was tested using gradient PCR. Relative expression was compared between genotypes and treatments using target primers and primers to the housekeeping gene *ACTIN7* for normalization (For primer sequences see, Data sheet 3). Agilent Brilliant III SYBR was used in conjunction with Agilent Aria MX qPCR machine and analysis performed using the ΔΔCT comparative quantification method (Livak and Schmittgen, 2001).

### rRNA analysis

Total RNA was extracted and quantified as described above and run on an Agilent Tapestation 2200. The concentration of RNA peaks of appropriate sizes corresponding to the abundant ribosomal RNAs (23S, 23Sb, 18S and 28S) were determined with provided software normalized to a RNA ladder standard. As no differences in cytosolic ribosome abundance was observed in the mutants, 18S rRNA was used to normalize chloroplast rRNA concentrations within samples to aid comparison between genotypes.

### Western blotting

The BIORAD Mini PROTEAN system was used for gel casting, running and transfer. 10% polyacrylamide gels (resolving gel: 0.38 M tris-HCl pH 8.8, 10% (w/v) acrylamide 0.1% (w/v) SDS, 0.05% (w/v) APS, 0.07% TEMED; stacking gel: 132 mM tris-HCl pH 6.8, 4% (w/v) acrylamide, 0.1% (w/v) SDS, 0.05% (w/v) APS, 0.15% (v/v) TEMED) were used to separate protein samples by gel electrophoresis. Separated proteins were transferred to PVDF membranes overnight at 4°C. Blotted membranes were blocked with 5% Marvel semi-skimmed milk in TBST for one hour. Membranes were probed with primary antibodies [anti-LHCA4, RubL, UGPase (Agrisera-AS01 008, AS03 037A, AS05 086), β-Glucuronidase (N-Terminal (Sigma-G5420), GFP (Roche-11814460001), Puromycin (Merk-MABE343), diluted 1:1000-1:5000 in TBST for three hours. Membranes washed three times for 5 minute in TBST. Membranes were probed with appropriate secondary antibodies, anti-Rabbit-Hrp, anti-Mouse-Hrp (Sigma A0545, A9917) diluted in TBST for one hour. Membranes were washed three times for 5 minutes in TBST. Membranes were incubated with Pierce™ ECL Western Blotting Substrate (Thermo Scientific-32106) for one minute. Membranes were exposed to X-ray film (FUJIFILM SUPER RX) in a HI-SPEED-X intensifying screen binder in a dark room. Films were developed with an Xograph Compact X4 Automated Processor and photographed on a light box with a Nikon D40 SLR camera.

### Chloroplast isolations

Chloroplasts were isolated from short day grown plants as (Kley et al., 2010). Two tubes containing 2g of fresh rosette leaf per sample were mixed with 23ml of chilled 1X isolation buffer each (0.6M sorbitol, 0.1M HEPES, 10mM EDTA, 10mM EGTA, 2mM MgCl_2_, 20mM NaHCO_3_ and 1mM DTT). Tissue was blended 3 times for 10 seconds with IKA T25 digital Ultra TURRAX at speed setting 3. Homogenate was poured into pre-wetted 38µm pore polyester mesh and filtered again through double layer of pre-wetted 22µm pore nylon cloth. Suspension was gently load onto two prepared falcon tubes containing equal volumes of 2X isolation buffer and Percoll™ (GE Healthcare-17-0891-02). Samples were centrifuged in a swing bucket centrifuge at 1200Xg for 10 minutes with brakes off. Upper layers were and pellet washed with 8ml 1X isolation buffer, inverting to mix. Samples were centrifuged again at 1000Xg for 5 minutes. Supernatants were removed using serological pipette and discarded. Pellets were resuspended in 5ml 1X isolation buffer loaded gently onto a 50% percoll isolation buffer. Samples were centrifuged at 1200Xg for 10 minutes with brakes off. Upper layers removed with a serological pipette. Pellets were washed with 10ml 1X isolation buffer, inverting to mix. Samples were centrifuged at 1000Xg for 5 and supernatant. Samples were centrifuged one final time at 1000Xg and remaining supernatant removed. Chloroplast pellets were frozen in liquid nitrogen. Proteins were solubilized in 2Xlaemmli buffer and quantified with the Biorad RC DC™ protein quantification assay.

For chloroplast puromycin incorporation assays chloroplast enrichment was undertaken as above with the following adjustments: DTT excluded from isolation buffer. After the second percoll gradient chloroplast enriched pellets were washed once in resuspension buffer (1.6M sorbitol, 0.1M HEPES, 2.5mM EDTA, 2.5mM EGTA, 0.5mM MgCl_2_, 20mM NaHCO_3_). Supernatants were removed and pellet resuspended in 2ml of fresh resuspension buffer. 0.4ml was taken for time-point 0 before adding 1.6µl of 50mM Puromycin dihydrochloride (Sigma-P8833), resuspensions were incubated in low light for 2 hours, 0.4ml was taken at each time-point and centrifuged at 12000Xg for 1 minute, supernatants removed and pellet snap frozen in liquid nitrogen. Protein extracts were prepared as above.

### Thylakoid composition analysis-Blue native gels and 2Ds

Thylakoids were isolated from 5-week-old plants grown in 8 h light/ 16 h dark at photosynthetic photon flux density (PPFD) of 100 μmol m^-2^ s^-1^, 50 % humidity, +23 °C, as described in (Koskela et al., 2018). Protein concentration was determined using BioRad’s DC Protein assay according to the manufacturer’s protocol.

Isolated thylakoids were diluted with ice-cold 25BTH20G [25 mM BisTris/HCl (pH 7.0), 20% (w/v) glycerol and 0.25 mg/ml Pefabloc, 10 mM NaF] buffer to a concentration of 10 mg protein/mL. An equal volume of detergent solution (β-dodecyl maltoside (DM) in BTH buffer) was added to a final concentration of 1% w/v. The membranes were solubilized 5 min on ice and the insolubilized material was removed by centrifugation at 18 000 x g at 4°C for 20 min. The supernatant was supplemented with Serva Blue G buffer [100 mM BisTris/HCl (pH 7.0), 5 M ACA, 30 %(w/v) sucrose and 50 mg/mL Serva Blue G] to introduce negative charge to the protein complexes. The Blue native (BN) gel (3.5-12.5% acrylamide) were prepared as described in Järvi et al 2011. The protein complexes from the sample were separated by the BN gel and the individual subunits further resolved with 2D-SDS-PAGE (12% acrylamide, 6 M urea) as in Järvi et al 2011). The proteins were visualized with SYPRO® Ruby staining according to Invitrogen Molecular Probes™ instructions, or with silver staining (Blum, 1987).

### Dual PAM measurements

Dual-PAM measurements were performed with Dual-PAM-100 (Heinz Walz GmbH) equipped with DUAL-E emitter and DUAL-DR detector units, using a red measuring beam for fluorescence and red actinic light. Simultaneously, the oxidation state of P700 was monitored by measuring the difference of the 875 nm and 830 nm transmittance signals. Prior to the measurements plants were dark-adapted for 30 min and F_0_, F_M_ and P_M_ were determined according to the Dual-PAM-100 protocol. For the light curve measurements the plants were subjected to illumination steps of 3 min at light intensities of 25-1000 μmol photons m^-2^ s^-1^ followed by a saturating flash (700 ms) to determine F_M’_ and P_M’_. For the induction curve, dark-adapted plants were illuminated with 1000 μmol photons m^-2^ s^-1^ actinic light and a saturating flash was applied every 2 min. The quantum yield of PSII (Y(II)) was calculated (Genty et al., 1989) from the fluorescence data, while the P700 signal was used for the quantum yield of PSI (Y(I)) (Klughammer and Schreiber, 2008). The relative rates of electron transfer through PSII and PSI, ETR(II) and ETR (I), respectively, were calculated as Y(II) x PPFD x 0.84 × 0.5 and Y(I) x PPFD x 0.84 × 0.5.

### Quantitative proteome analysis

Col-0 and *not4a* plants were grown on 1/2 MS medium, supplemented with 0.8% sugar, for 30d under short day conditions. Plant material from three independent biological replicates was homogenized in Rensink extraction buffer (50 mM Tris/HCl pH 7.5, 100 mM NaCl, 0.5% (v/v) TritonX-100, 2 mM DTT and protease inhibitor cocktail (Sigma-Aldrich)). LC separation and HD-MSE data acquisition was performed as previously described (Helm et al., 2014). In short, 100µg protein were digested in solution with RapiGestTM with 1 µg Trypsin (Promega) over night. Peptide pellets were dissolved in 2% (v/v) ACN, 0.1% (v/v) FA, and subjected to LC on an ACQUITY UPLC System coupled to a Synapt G2-S mass spectrometer (Waters, Eschborn, Germany). For quantification, the sample was spiked with 10 fmol rabbit glycogen phosphorylase and the abundance of the three most intense peptide ions were used as s reference value for 10 fmol (Hi-3 method, Silva et al., 2006). Data analysis and quantification was carried out by ProteinLynx Global Server (PLGS 3.0.1, Apex3D algorithm v. 2.128.5.0, 64 bit, Waters, Eschborn, Germany) with automated determination of chromatographic peak width as well as MS TOF resolution. Database query was as follows: Peptide and fragment tolerances were set to automatic, two fragment ion matches per peptide, five fragment ions for protein identification, and two peptides per protein were required for identification. Primary digest reagent was trypsin with one missed cleavage allowed. The false discovery rate (FDR) was set to 4% at the protein level. MSE data were searched against the modified A. thaliana database (TAIR10, ftp://ftp.arabidopsis.org) containing common contaminants such as keratin (ftp://ftp.thegpm.org/fasta/cRAP/crap.fasta). All quantitative proteomics data were obtained with three independent biological replicates that were measured in three technical replicates each, giving rise to nine measurements per sample. Mapman was used to assign proteins to functional groups(Thimm et al., 2004). Statistical testing was based on a two-sided T-test (Helm et al., 2014).

### Photoinhibition assays

Detached leaves of mature plants grown under standard conditions for five weeks were floated on water or 2.3 mM lincomycin for 16 h in darkness after which they were illuminated under with high light of 1000 μmol photons m^-2^ s^-1^ for 1.5 h. Thereafter, the water samples were moved to standard growth conditions for recovery. PSII efficiency (F_V_/F_M_) was measured after a 20 min dark-adaptation using FluorPen FP 110 (Photon Systems Instruments). The amount of D1 protein was measured by western blotting with protein samples collected at indicated timepoints. Total protein samples were isolated by homogenizing frozen leaf material in 100 mM Tris-HCl pH 8.0, 50 mM Na-EDTA, 0.25 mM NaCl, 0.75 (w/v) % SDS, 1 mM DTT followed by 10 min incubation at 68 °C. The extracts were clarified by centrifugation at 12 000 x g for 10 min. Protein concentration was determined using BioRad’s DC Protein assay according to the manufacturer’s protocol. Solubilized protein samples were separated by SDS-PAGE (12% acrylamide, 6 M urea) and immunoblotted with rabbit D1 DE-loop antibody (Kettunen et al., 1996) used as a 1:8000 dilution. LI-COR Goat anti-rabbit IRDye® 800CW 2nd antibody was used for detection according to manufacturer’s instructions.

### Confocal Microscopy

4 day old pPGR3::PGR3-YFP expressing seedlings were imaged with a Nikon Ti microscope connected to an A1R confocal system equipped with a Plan Apochromat 60x/1.2 WI DIC H lens, with 514nm and 637nm laser lines, to image YFP and chlorophyll auto-fluorescence respectively. Laser power, gain and pinhole settings used were identical between lines, YFP was imaged prior to chloroplast auto-fluorescence and settings compared to WT seedlings as a YFP negative control. At least 4 seedlings per genetic background were imaged.

### Yeast-two hybrid

PGR3 and NOT4A coding sequences were cloned into Invitrogen pENTR™/D-TOPO™ (ThermoFisher-K240020) and sequenced before ligation into pGADCg and pGBKCg respectively (For primer sequences see, Data sheet 3). Matchmaker (AH109) yeast was transformed with 1µg pGADCg and pGBKCg vectors containing NOT4A, PGR3 or empty vectors as described, in 100µl TB buffer (2:1 50% PEG 3350MW, 1M Lithium Acetate, 0.6% ß-mercaptoethanol). 37°C for 45 minutes and then spread on agar plates containing yeast Nitrogen Base without Amino acids supplemented with synthetic amino acid Drop out (DO) - leu-trp agar (Formedium-CYN0401, DSCK172) and grown at 30°C for 2-3 days before resuspending single colonies in 100µl sterile water and transferring 20µl to DO -leu-trp-his-ade and DO -leu-trp agar plates (Formedium-DSCK272, DSCK172). Plates were grown at 30°C for 2-3 days before photographing with Nikon D40 SLR camera.

## Acknowledgments

This work was supported by Biotechnology and Biological Sciences Research Council grants [BB/M020568/1 and BB/T004002/1] and a European Research Council grant [ERC Starting Grant 715441-GasPlaNt] to D.J.G, an EMBO Short-Term Fellowship [Grant number 8104] to M.B, Academy of Finland grants [307335 and 321616] for AI, MR and PM, and the Doctoral Programme in Molecular Life Sciences at the University of Turku (AI). SB gratefully acknowledges support from the DFG with grant number BA 1902/3-2. We thank Dr Alessandro Di Mio and the BALM facility at the University of Birmingham for support with confocal analyses.

## Author contributions

M.B. and D.J.G. conceived and designed the overall project, and A.I and P.R. designed the photosynthetic analyses. M.B., A.I., O.A., A.C.P., R.E., A-M.L., R.O., M.R. and D.J.G performed experiments. J.G and S.B. performed the quantitative proteome analysis. M.B., A.I., P.M. and D.J.G. analysed data. M.B and D.J.G wrote the manuscript with input from all authors.

## Supplemental information

Supplemental figures 1-7

Supplemental data file 1-Transcriptome profiling

Supplemental data file 2-Proteome profiling

Supplemental data file 3-Primer list

