## Supplemental Figures for "The *Arabidopsis* NOT4A E3 ligase coordinates PGR3 expression to regulate chloroplast protein translation"

### Slide 1
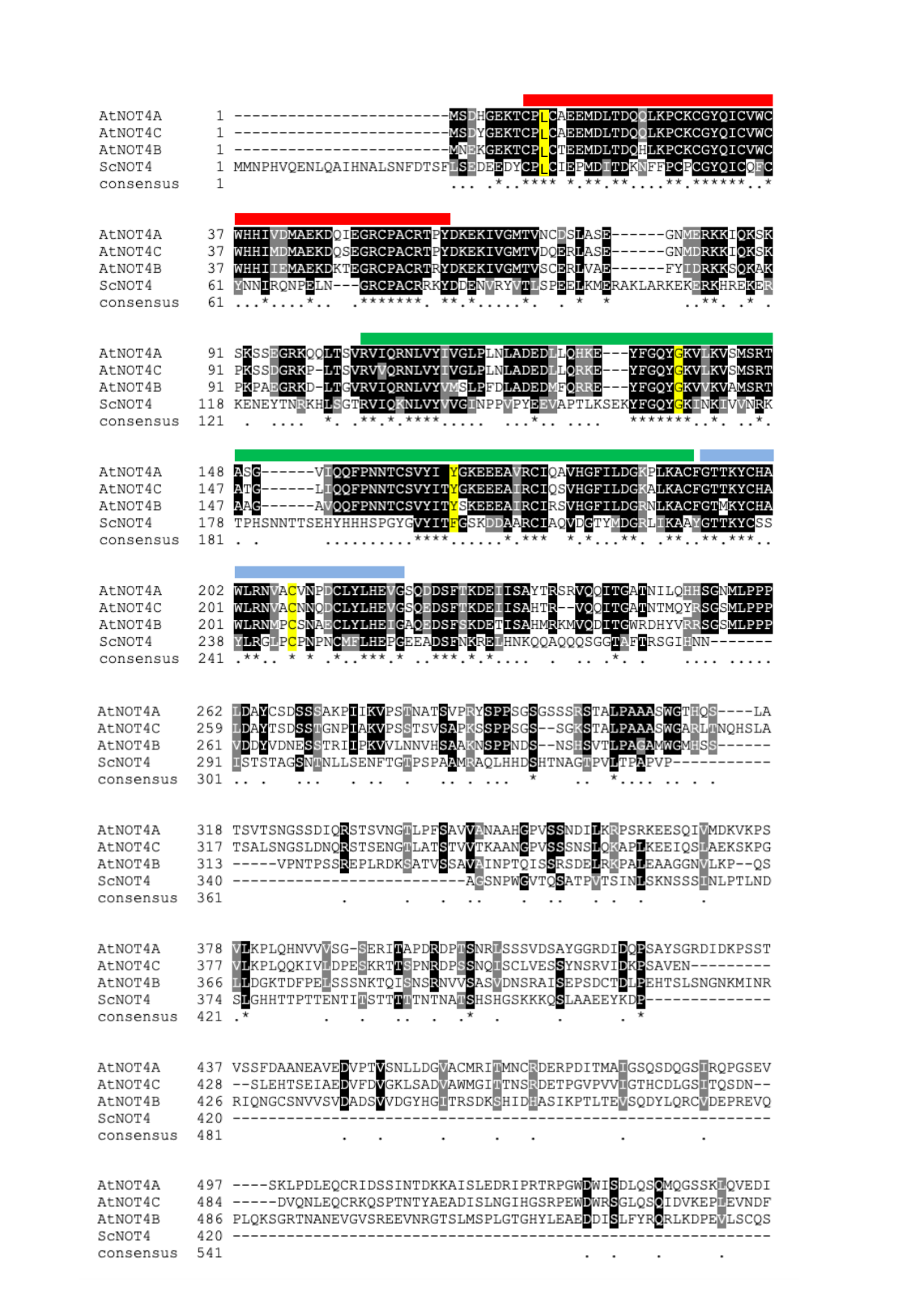

L
L
L
L

### Slide 2
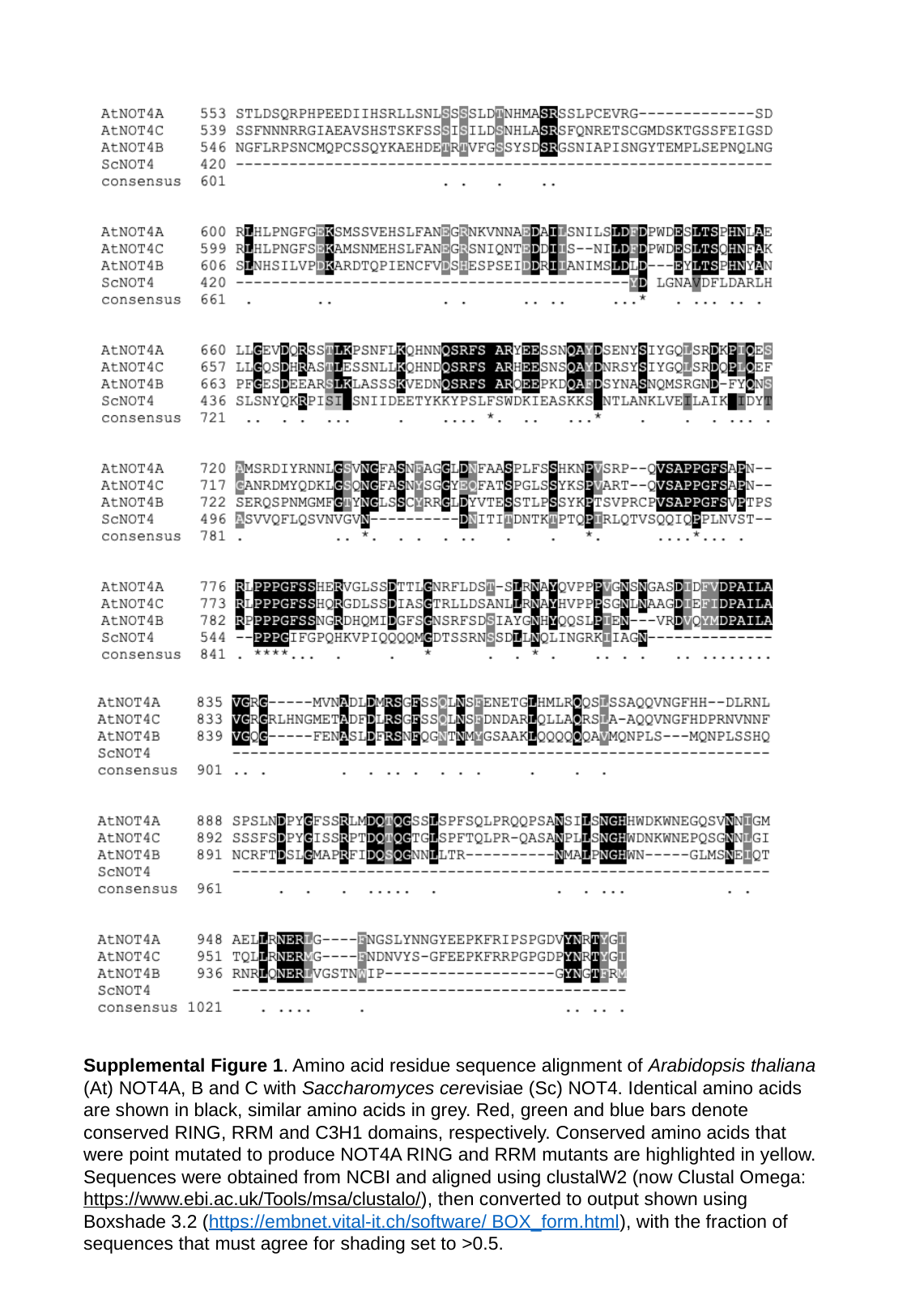

Supplemental Figure 1. Amino acid residue sequence alignment of Arabidopsis thaliana (At) NOT4A, B and C with Saccharomyces cerevisiae (Sc) NOT4. Identical amino acids are shown in black, similar amino acids in grey. Red, green and blue bars denote conserved RING, RRM and C3H1 domains, respectively. Conserved amino acids that were point mutated to produce NOT4A RING and RRM mutants are highlighted in yellow. Sequences were obtained from NCBI and aligned using clustalW2 (now Clustal Omega: https://www.ebi.ac.uk/Tools/msa/clustalo/), then converted to output shown using Boxshade 3.2 (https://embnet.vital-it.ch/software/ BOX_form.html), with the fraction of sequences that must agree for shading set to >0.5.

### Slide 3
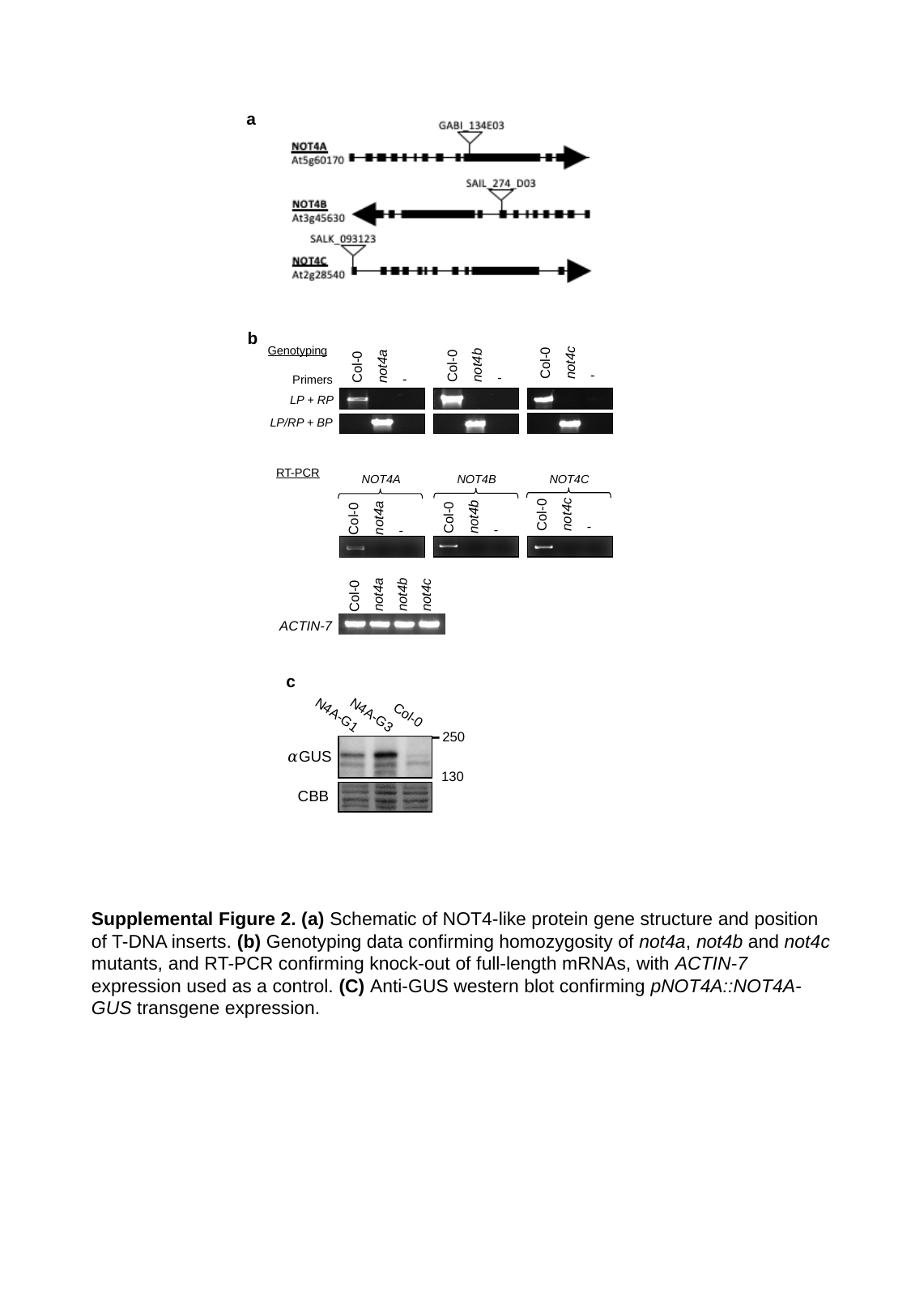

a
b
Genotyping
Col-0
not4c
not4b
Col-0
Col-0
not4a
-
-
-
Primers
LP + RP
LP/RP + BP
RT-PCR
NOT4A
NOT4B
NOT4C
Col-0
not4c
not4b
Col-0
Col-0
not4a
-
-
-
not4b
not4c
not4a
Col-0
ACTIN-7
c
N4A-G1
N4A-G3
Col-0
-
250
𝛼GUS
130
CBB
Supplemental Figure 2. (a) Schematic of NOT4-like protein gene structure and position of T-DNA inserts. (b) Genotyping data confirming homozygosity of not4a, not4b and not4c mutants, and RT-PCR confirming knock-out of full-length mRNAs, with ACTIN-7 expression used as a control. (C) Anti-GUS western blot confirming pNOT4A::NOT4A-GUS transgene expression.

### Slide 4
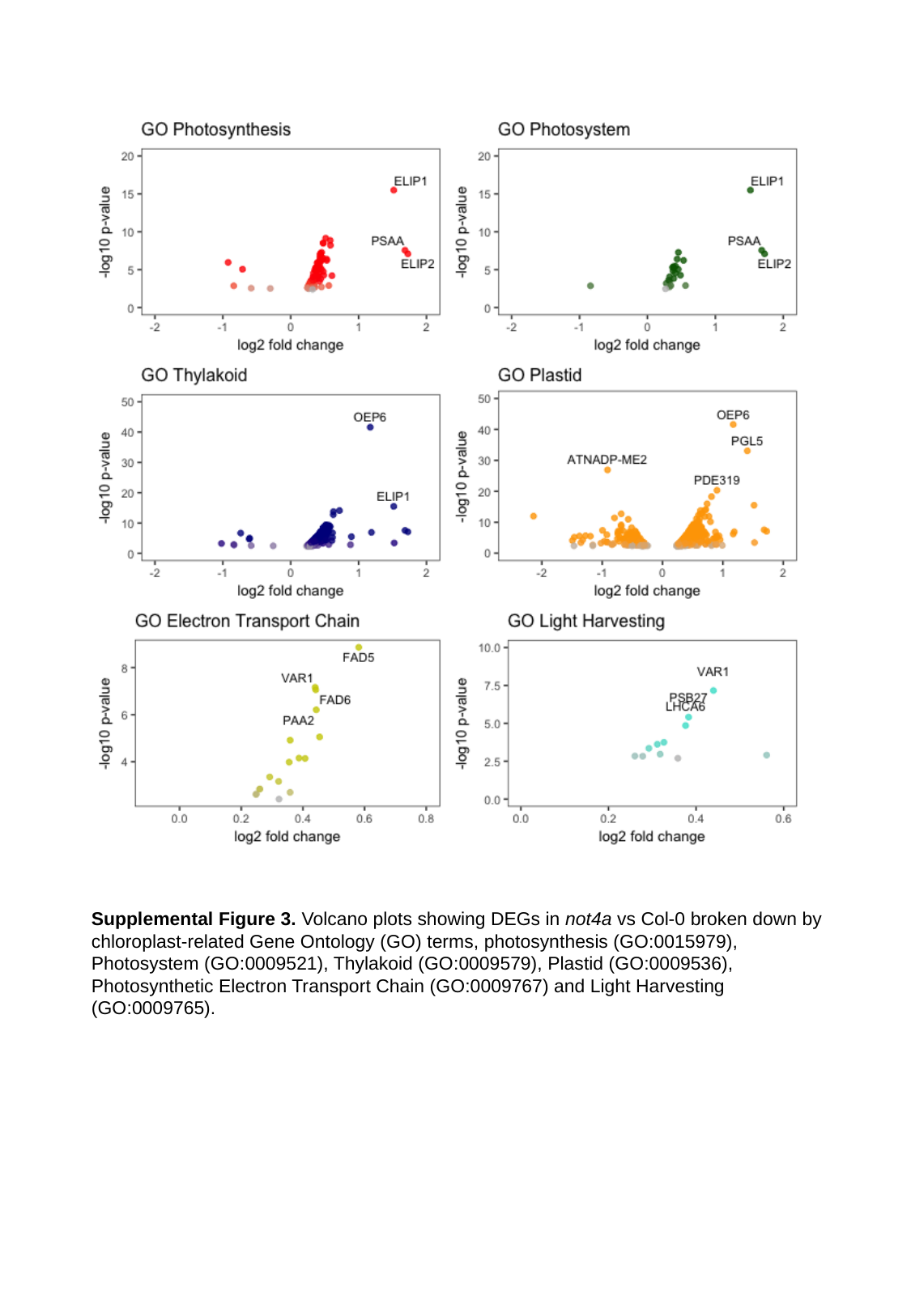

Supplemental Figure 3. Volcano plots showing DEGs in not4a vs Col-0 broken down by chloroplast-related Gene Ontology (GO) terms, photosynthesis (GO:0015979), Photosystem (GO:0009521), Thylakoid (GO:0009579), Plastid (GO:0009536), Photosynthetic Electron Transport Chain (GO:0009767) and Light Harvesting (GO:0009765).

### Slide 5
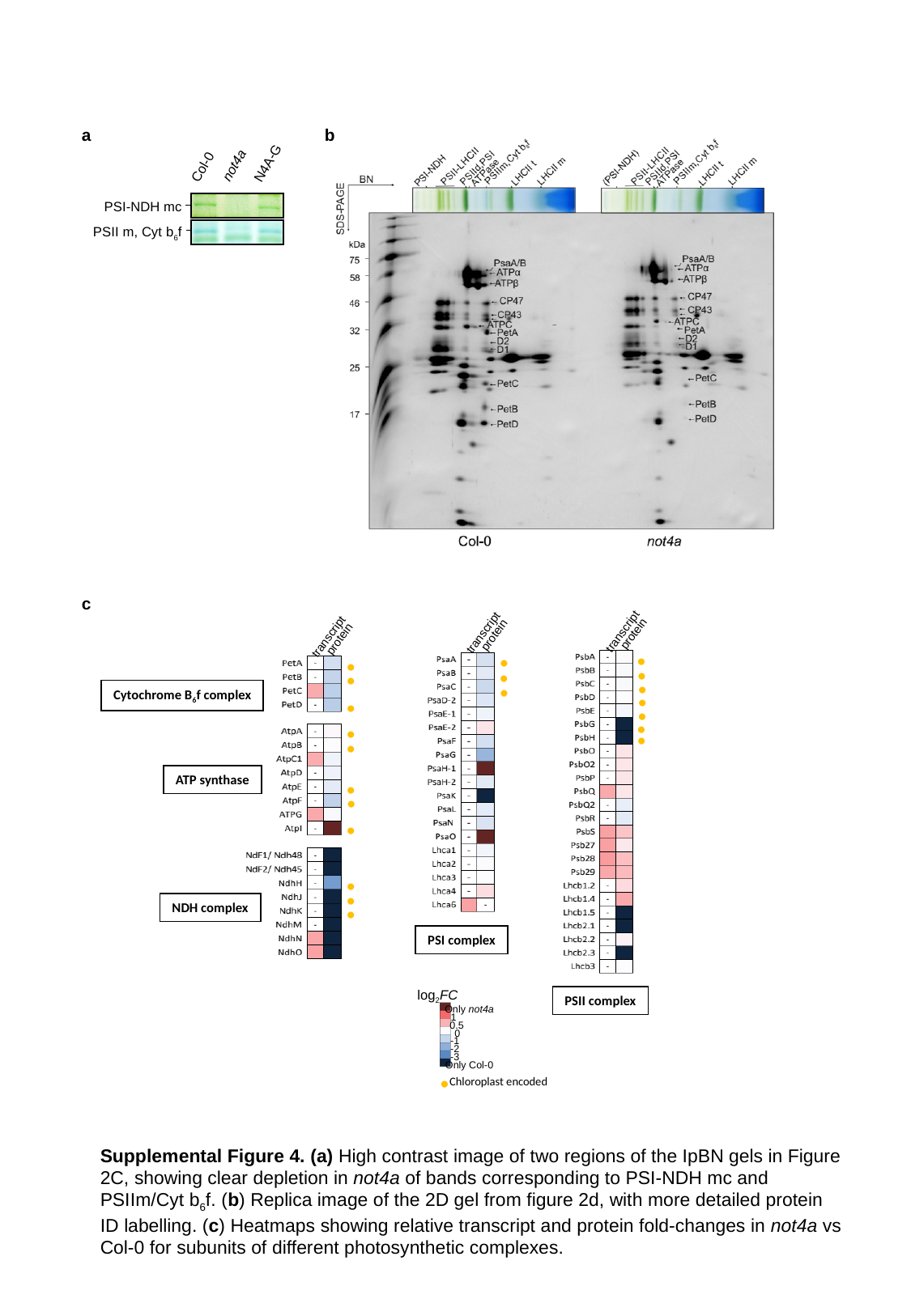

a
b
not4a
Col-0
PSI-NDH mc
PSII m, Cyt b_6_f
N4A-G
c
transcript
protein
transcript
protein
transcript
protein
Cytochrome B_6_f complex
ATP synthase
NDH complex
PSI complex
log_2_FC
PSII complex
Only not4a
1
0.5
0
-1
-2
-3
Only Col-0
•
•
•
•
•
•
•
•
•
•
•
•
•
•
•
•
•
•
•
•
•
•
Chloroplast encoded
Supplemental Figure 4. (a) High contrast image of two regions of the IpBN gels in Figure 2C, showing clear depletion in not4a of bands corresponding to PSI-NDH mc and PSIIm/Cyt b_6_f. (b) Replica image of the 2D gel from figure 2d, with more detailed protein ID labelling. (c) Heatmaps showing relative transcript and protein fold-changes in not4a vs Col-0 for subunits of different photosynthetic complexes.

### Slide 6
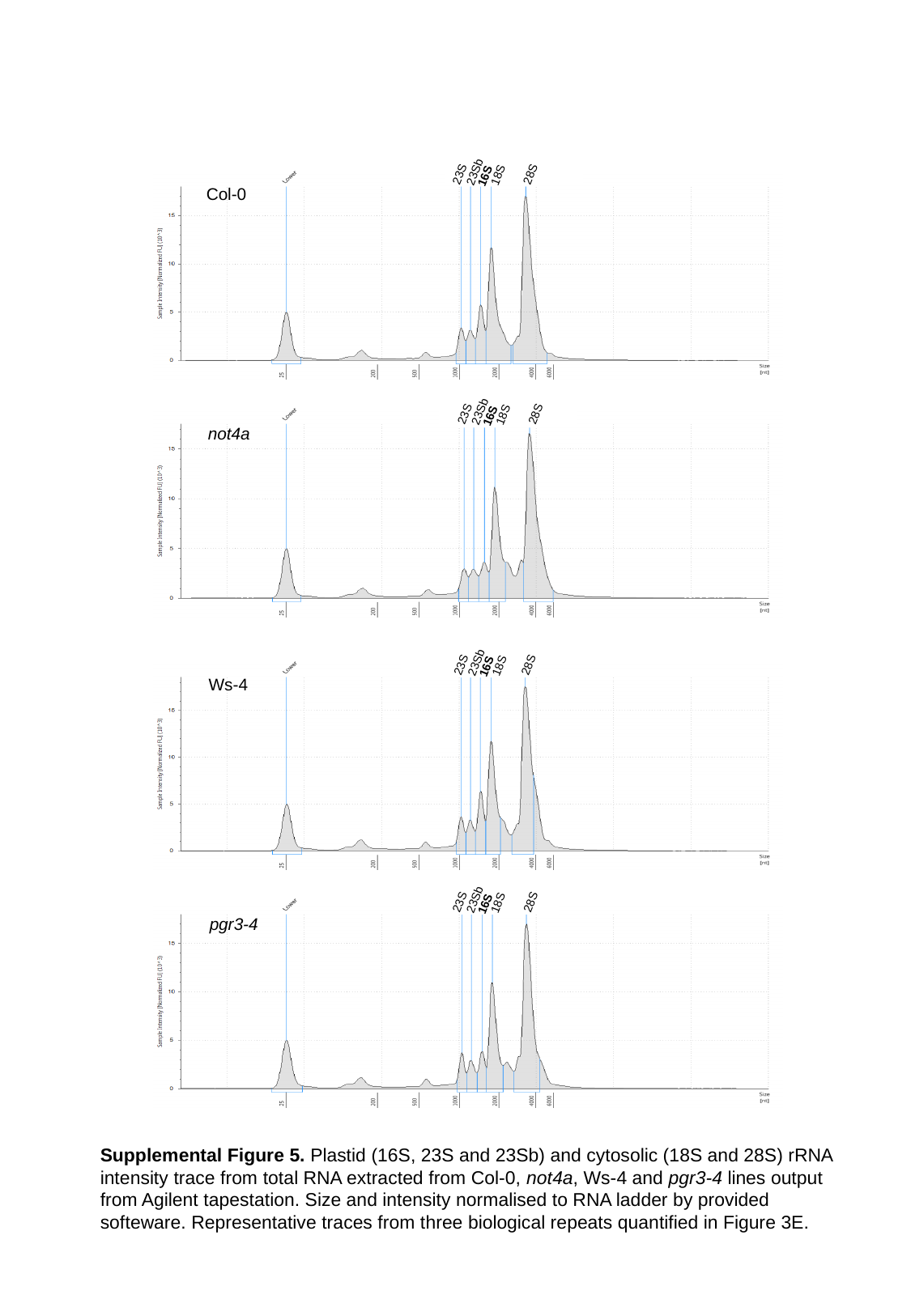

23Sb
23S
28S
18S
16S
Col-0
23Sb
23S
28S
18S
16S
not4a
23Sb
23S
28S
18S
16S
Ws-4
23Sb
23S
28S
18S
16S
pgr3-4
Supplemental Figure 5. Plastid (16S, 23S and 23Sb) and cytosolic (18S and 28S) rRNA intensity trace from total RNA extracted from Col-0, not4a, Ws-4 and pgr3-4 lines output from Agilent tapestation. Size and intensity normalised to RNA ladder by provided softeware. Representative traces from three biological repeats quantified in Figure 3E.

### Slide 7
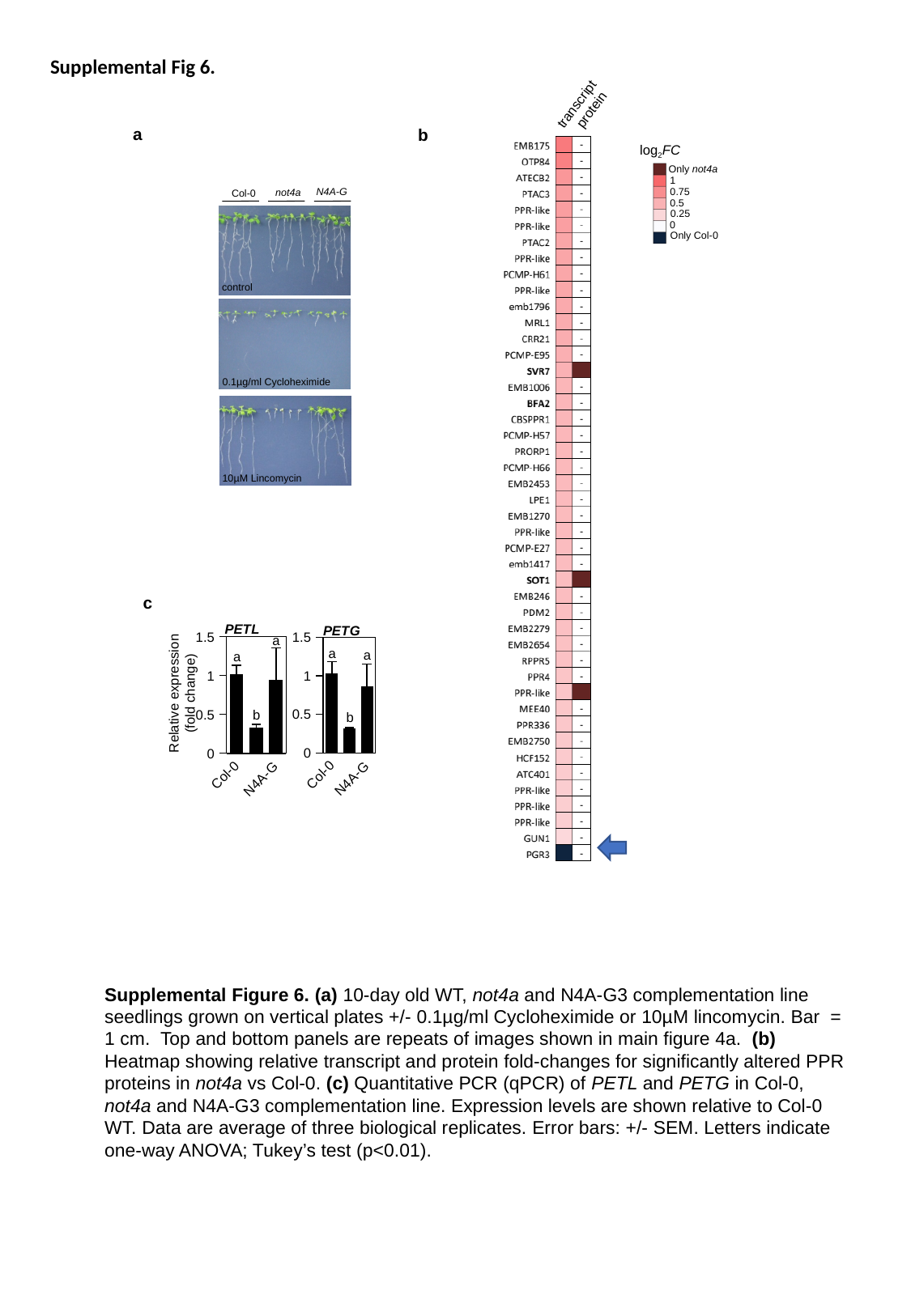

Supplemental Fig 6.
transcript
protein
a
b
log_2_FC
Only not4a
1
0.75
0.5
0.25
0
Only Col-0
 not4a
 Col-0
 N4A-G
control
0.1µg/ml Cycloheximide
10µM Lincomycin
#### Chart: PETG
| Category | PETG |
|---|---|
| Col-0 | 1.026437122658245 |
| not4a | 0.318404858741099 |
| N4A-G | 0.85912153074681 |a
a
a
a
b
b
#### Chart: PETL
| Category | PETL |
|---|---|
| Col-0 | 1.014980197650193 |
| not4a | 0.326142296765717 |
| N4A-G | 0.94164034243861 |c
Relative expression
(fold change)
Supplemental Figure 6. (a) 10-day old WT, not4a and N4A-G3 complementation line seedlings grown on vertical plates +/- 0.1µg/ml Cycloheximide or 10µM lincomycin. Bar = 1 cm. Top and bottom panels are repeats of images shown in main figure 4a. (b) Heatmap showing relative transcript and protein fold-changes for significantly altered PPR proteins in not4a vs Col-0. (c) Quantitative PCR (qPCR) of PETL and PETG in Col-0, not4a and N4A-G3 complementation line. Expression levels are shown relative to Col-0 WT. Data are average of three biological replicates. Error bars: +/- SEM. Letters indicate one-way ANOVA; Tukey’s test (p<0.01).

### Slide 8
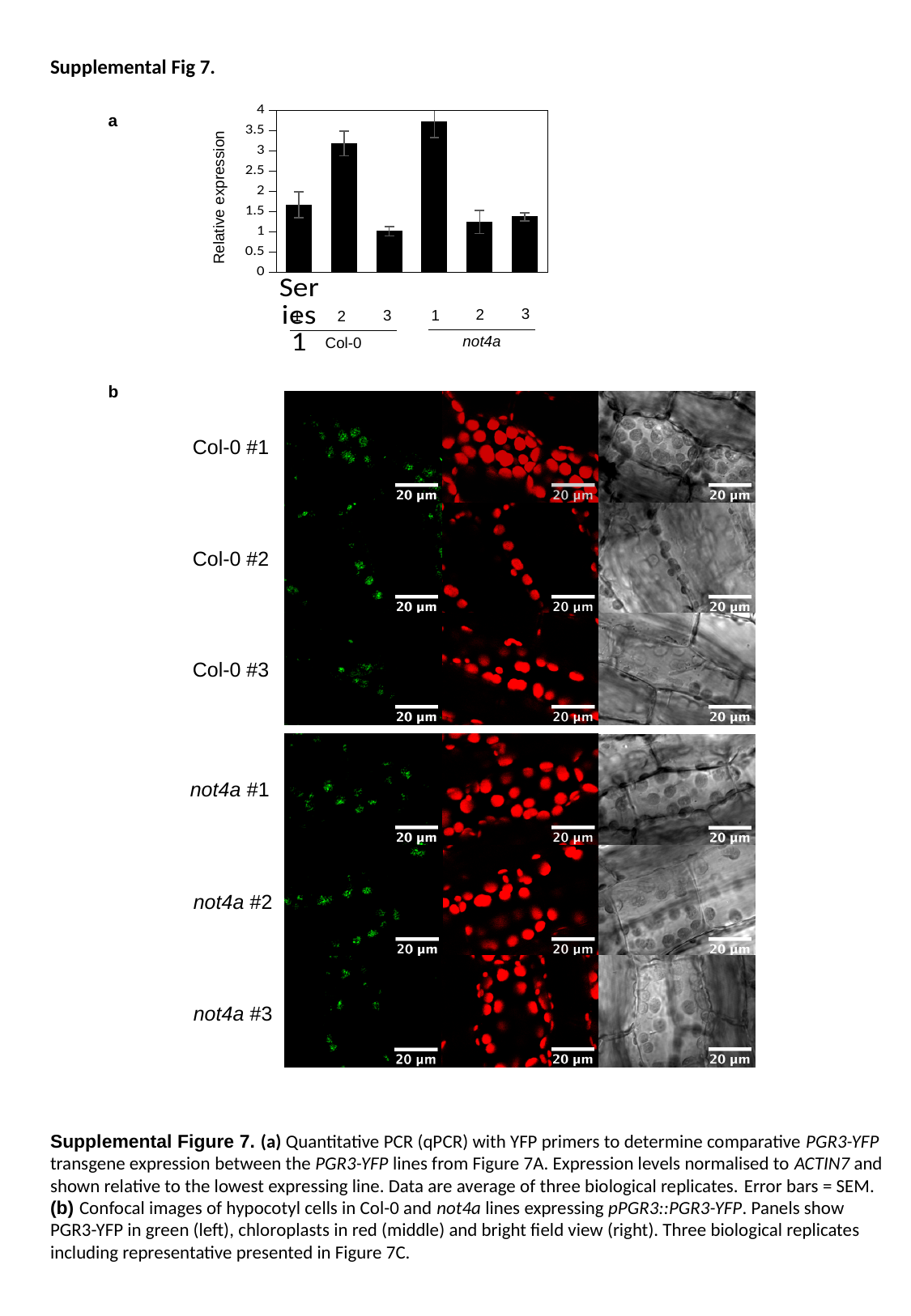

Supplemental Fig 7.
#### Chart
| Category | |
|---|---|
| | 1.667742570577751 |
| | 3.190302394083522 |
| | 1.01326996757685 |
| | 3.717978569380043 |
| | 1.243315283733672 |
| | 1.371593504412105 |a
Relative expression
3
2
3
1
2
1
not4a
Col-0
b
Col-0 #1
Col-0 #2
Col-0 #3
not4a #1
not4a #2
not4a #3
Supplemental Figure 7. (a) Quantitative PCR (qPCR) with YFP primers to determine comparative PGR3-YFP transgene expression between the PGR3-YFP lines from Figure 7A. Expression levels normalised to ACTIN7 and shown relative to the lowest expressing line. Data are average of three biological replicates. Error bars = SEM. (b) Confocal images of hypocotyl cells in Col-0 and not4a lines expressing pPGR3::PGR3-YFP. Panels show PGR3-YFP in green (left), chloroplasts in red (middle) and bright field view (right). Three biological replicates including representative presented in Figure 7C.
